# Discovery and reduction of nonspecific activities of the major herbicide-resistance gene BAR

**DOI:** 10.1101/122184

**Authors:** Bastien Christ, Ramon Hochstrasser, Luzia Guyer, Rita Francisco, Sylvain Aubry, Stefan Hörtensteiner, Jing-Ke Weng

## Abstract

Herbicide resistance is a major trait of genetically modified (GM) crops. Currently, resistance to phosphinothricin (also known as glufosinate) is the second most widespread genetically engineered herbicide-resistance trait in crops after glyphosate resistance^1,2^. Resistance to phosphinothricin in plants is achieved by transgenic expression of the bialaphos resistance (*BAR*) or phosphinothricin acetyltransferase (*PAT*) genes, which were initially isolated from the natural herbicide bialaphos-producing soil bacteria *Streptomyces hygroscopicus* and *S. viridochromogenes*, respectively^3,4^. Mechanistically, *BAR* and *PAT* encode phosphinothricin acetyltransferase, which transfers an acetyl group from acetyl coenzyme A (acetyl-CoA) to the α-NH_2_ group of phosphinothricin, resulting in herbicide inactivation^1^. Although early in vitro enzyme assays showed that recombinant BAR and PAT exhibit substrate preference toward phosphinothricin over the 20 proteinogenic amino acids^1^, whether transgenic expression of BAR and PAT affects plant endogenous metabolism in vivo was not known. Combining metabolomics, plant genetics, and biochemical approaches, we show that transgenic BAR indeed converts two plant endogenous amino acids, aminoadipate and tryptophan, to their respective N-acetylated products in several plant species examined. We report the crystal structures of BAR, and further delineate structural basis for its substrate selectivity and catalytic mechanism. Through structure-guided protein engineering, we generated several BAR variants that display significantly reduced nonspecific activities compared to its wild-type counterpart. Our results demonstrate that transgenic expression of enzymes as a common strategy in modern biotechnology may render unintended metabolic consequences arisen from enzyme promiscuity. Understanding of such phenomena at the mechanistic level will facilitate better design of maximally insulated systems featuring heterologously expressed enzymes.

---

Phosphinothricin is a naturally occurring herbicide derived from the tripeptide antibiotic bialaphos produced by species of *Streptomyces* soil bacteria. Phosphinothricin is a structural analog of glutamate, and thereby inhibits glutamine synthetase, an essential enzyme for glutamine synthesis and ammonia detoxification in plants, giving rise to its herbicidal activity^3^. In the 1980s, the bialaphos resistance (*BAR*) gene and its closely related homolog phosphinothricin acetyltransferase (*PAT*) gene were isolated from *S. hygroscopicus* and *S. viridochromogenes*, respectively, and were later broadly used as transgenes to confer herbicide resistance in a variety of major genetically modified (GM) crops, including corn, soybean, canola, and cotton. In addition, *BAR* and *PAT* have also gained much utility in basic research as selection markers for generating transgenic plants^1^. Despite the prevalent use of *BAR* and *PAT* in the context of generating herbicide-resistant transgenic plants, whether these bacteria-derived enzymes may possibly interfere with plant endogenous metabolism has not been rigorously investigated.

In research not initially intended to address this issue regarding phosphinothricin-resistance trait, we carried out untargeted metabolomics analysis on senescent leaf extracts prepared from the *Arabidopsis thaliana clh2-1* mutant (FLAG_76H05), which contains a transfer DNA (T-DNA) insertion that abolishes the *CHLOROPHYLLASE 2* gene. This analysis revealed two metabolites that were ectopically accumulated at high levels in *clh2-1* compared to wild type (**Fig. 1a**). Using liquid chromatography-tandem mass spectrometry (LC-MS^2^), we identified these two metabolites as N-acetyl-L-aminoadipate and N-acetyl-L-tryptophan (referred to as acetyl-aminoadipate and acetyl-tryptophan, respectively, hereafter; **Fig. 1a** and **Supplementary Fig. 1**). Because the deficiency of CHLOROPHYLLASE 2, a serine esterase^5^, in *clh2-1* does not explain the accumulation of these acetylated metabolites, we hypothesized that the *BAR* gene present on the T-DNA as a selection marker in *clh2-1* might be responsible for their formation. To test this, we extended our metabolomics analysis to additional Arabidopsis T-DNA insertional mutants unrelated to chlorophyll metabolism that carry either *BAR* (e.g. mutants from the FLAG^6^ and SAIL^7^ collections) or alternative antibiotic selection markers (e.g. mutants from the SALK^8^ and GABI^9^ collections). Senescent leaves of all six T-DNA mutants carrying *BAR* manifested accumulation of acetyl-aminoadipate and acetyl-tryptophan, while these metabolites were significantly lower or not detected in wild-type plants and T-DNA mutants containing alternative selection markers (**Fig. 1b**). These results indicate that the ectopic accumulation of these metabolites is likely resulted from the nonspecific N-acetyltransferase activities of transgenic BAR acting upon plant endogenous amino acids.

**Figure 1.**
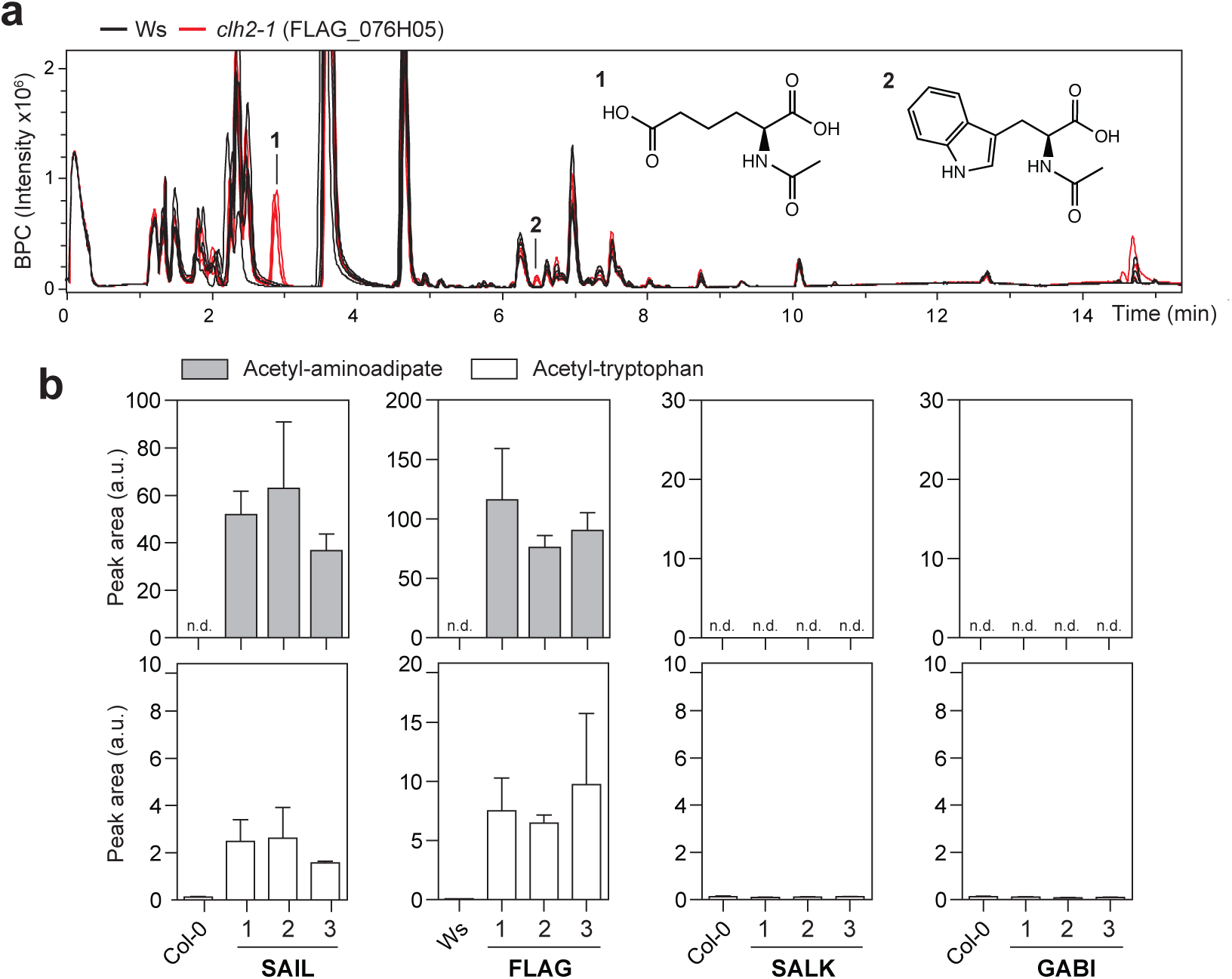
Accumulation of acetyl-aminoadipate and acetyl-tryptophan in senescent leaves of Arabidopsis carrying the BAR transgene. (**a**) Metabolite profiles of senescent leaves from Wassilewskija (Ws) and *clh2-1* (FLAG_076H05), displayed as base peak chromatograms (BPC), reveal the ectopic accumulation of acetyl-aminoadipate (1) and acetyl-tryptophan (2). BPC traces of four biological replicates are displayed. (**b**) Relative quantification of acetyl-aminoadipate and acetyl-tryptophan in Arabidopsis mutants from different insertional mutant collections that contain either BAR (SAIL and FLAG) or alternative selection marker genes (SALK and GABI). Error bars, mean ± s.d. (n = 4 biological replicates). This experiment was repeated at least three times with similar results. See **Supplementary Table 1** for absolute quantification. a.u., arbitrary unit; FW, fresh weight; n.d., not detected

The concentration of free tryptophan is low in photosynthetically active leaves, but increases significantly in senescent leaves^10^. This is due to enhanced proteolysis during senescence, facilitating remobilization of protein-bound nitrogen and other nutrients to sink organs, such as seeds^11^. Aminoadipate, an intermediate of lysine degradation, also exhibits a similar accumulation pattern during leaf senescence^12^. To test whether the BAR-catalyzed production of acetyl-aminoadipate depends on lysine degradation, we analyzed an Arabidopsis mutant from the FLAG collection, FLAG*_lkrsdh*, in which the *BAR*-containing T-DNA disrupts *At4g33150* encoding the Arabidopsis bifunctional lysine-ketoglutarate reductase/saccharopine dehydrogenase (LKR/SDH)^13^. LKR/SDH catalyzes the first committed step of lysine degradation, and, together with the subsequent aminoadipate semialdehyde dehydrogenase (AADH), converts lysine to aminoadipate (**Fig. 2a**). In a segregating population for the FLAG*_lkrsdh* locus, heterozygous, homozygous and wild-type plants were identified by genotyping, and subjected to LC-MS metabolic profiling after senescence induction (**Fig. 2b**). Acetyl-aminoadipate occurred at the highest level in the heterozygous mutant, but was greatly reduced in the homozygous mutant, suggesting that the ectopic accumulation of acetyl-aminoadipate in *BAR*-containing plants is dependent on LKR/SDH of the lysine degradation pathway in senescent leaves (**Fig. 2a**). By contrast, the relative abundance of acetyl-tryptophan in the segregating population of FLAG *lkrsdh* generally reflected the copy number of the BAR-containing T-DNA transgene, with approximately 2-fold acetyl-tryptophan level observed in the homozygotes compared to the heterozygotes (**Fig. 2b**). Furthermore, acetyl-aminoadipate and acetyl-tryptophan levels were approximately 10-20 fold higher in senescent leaves than those in green leaves (**Fig. 2b**), which is likely due to the increased availability of the corresponding free amino acids during senescence. Consistent with these observations in leaves, ectopic accumulation of acetyl-aminoadipate and acetyl-tryptophan was also observed in seeds of multiple *BAR*-containing T-DNA mutant lines compared to the wild-type controls (**Supplementary Fig. 2**). We quantified the absolute concentrations of acetyl-aminoadipate and acetyl-tryptophan in various tissues of *BAR*-containing transgenic Arabidopsis to range from 43 to 308 μg/g and from 0.26 to 28 μg/g fresh weight, respectively (**Supplementary Table 1**). These concentrations are substantial given that the total pool of free amino acids in senescent leaves of Arabidopsis is reportedly around 1000 μg/g fresh weight^14^.

**Figure 2.**
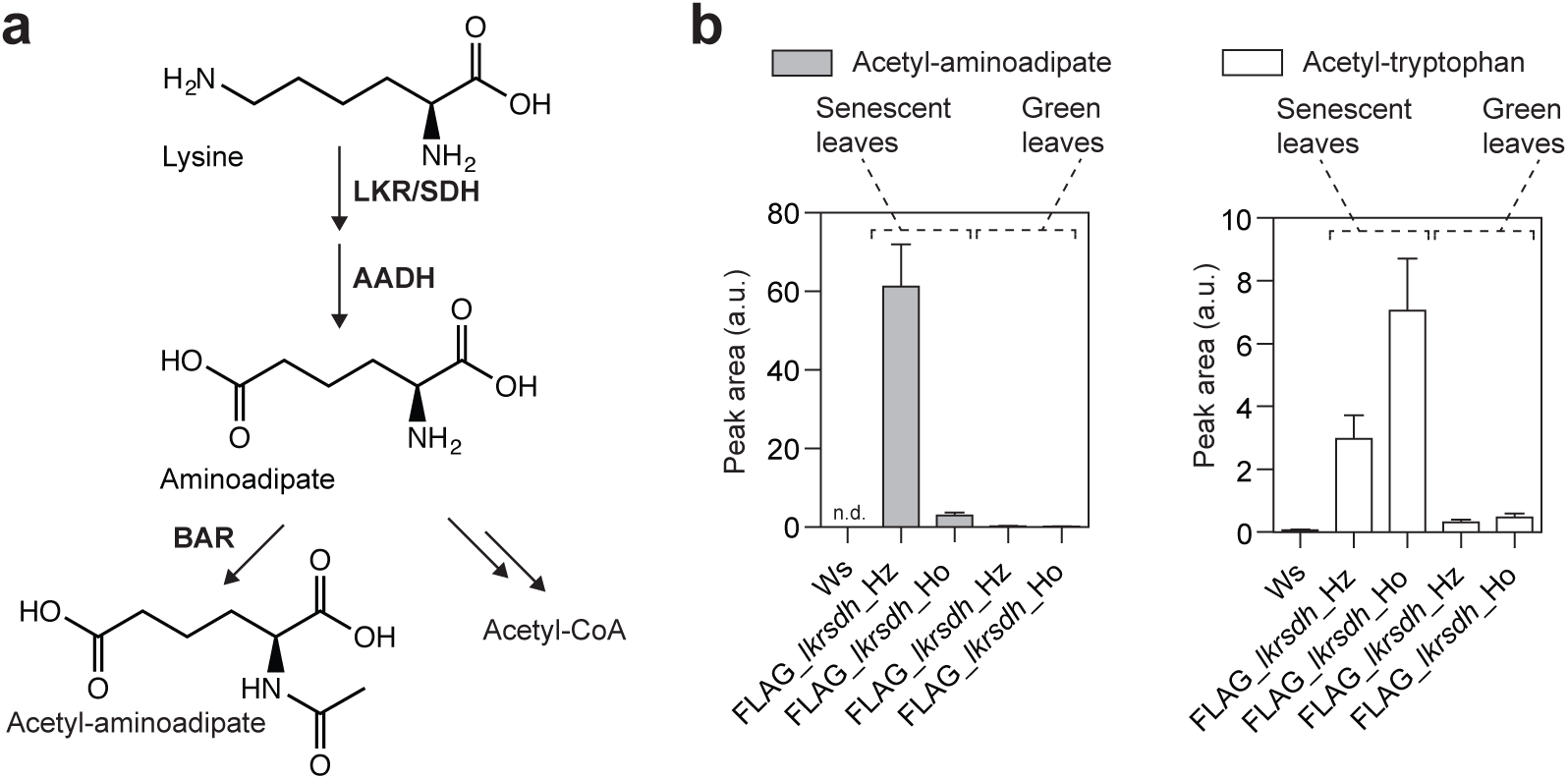
BAR-dependent accumulation of acetyl-aminoadipate and acetyl-tryptophan is linked to nitrogen remobilization during senescence. (**a**) Aminoadipate is derived from the lysine degradation pathway in plants, which can be metabolized by BAR as a nonspecific substrate. (**b**) Relative quantification of acetyl-aminoadipate and acetyl-tryptophan in green and senescent leaves from the heterozygous (Hz) and homozygous (Ho) *FLAG_lkrsdh* mutant, harboring a BAR-containing T-DNA that abolishes the Arabidopsis *LKR/SDH* gene. Error bars, mean ± s.d. (n = 3 biological replicates). a.u., arbitrary unit; n.d., not detected; Ws, Wassilewskija wild-type plants.

To assess whether the nonspecific activities of transgenic BAR also manifest in other plant hosts, we performed metabolic profiling of various tissue samples from phosphinothricin-resistant soybean (*Glycine max*) and Chinese mustard (*Brassica juncea*). Substantially increased accumulation of acetyl-aminoadipate and acetyl-tryptophan was also detected in some tissues of these transgenic crops (**Supplementary Fig. 3**), indicating that our findings regarding the in vivo nonspecific activities of BAR may apply broadly to a wide range of *BAR*-containing transgenic plants.

To shed light on the kinetic properties of BAR, we carried out pseudo-first-order enzyme kinetic assays using recombinant BAR against several native and non-native amino acid substrates (**Fig. 3** and **Supplementary Fig. 4**). Similar to published data^1,3,15^, N-acetylation of phosphinothricin exhibits Michaelis-Menten kinetics with an apparent K_M_ of approximately 132 μM (**Fig. 3**). Although BAR clearly showed N-acetyltransferase activities toward aminoadipate and tryptophan, kinetic constants for these non-native substrates could not be established, as both substrates reached solubility limit before reaching saturation concentration for BAR. Based on these results, we estimate that the *K*_M_ values of BAR against aminoadipate and tryptophan are at least in the millimolar range. BAR also exhibited relatively higher catalytic activity toward aminoadipate than tryptophan in vitro (**Fig. 3**).

**Figure 3.**
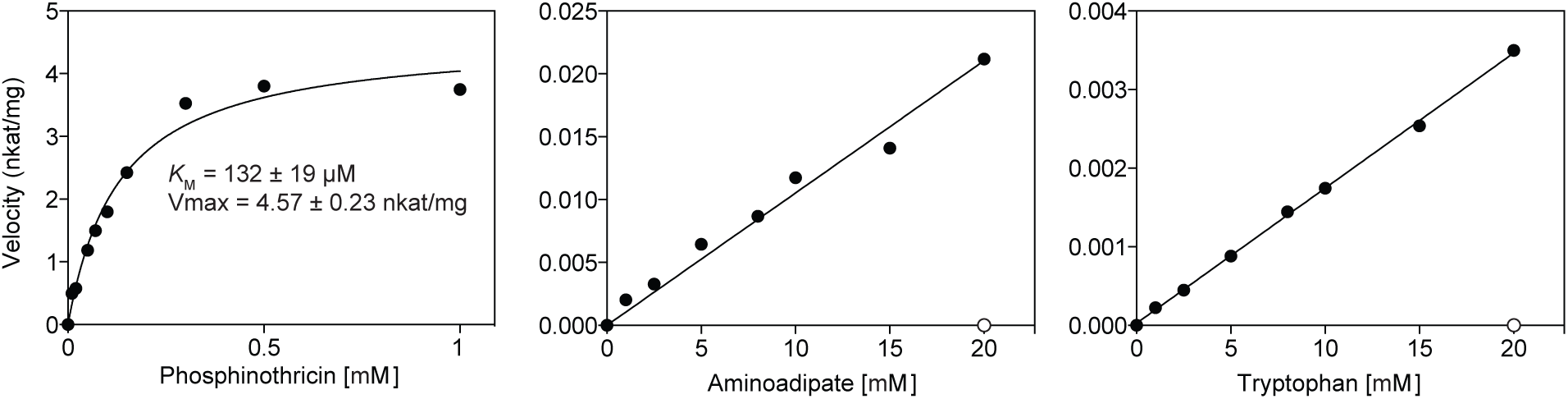
In vitro enzyme kinetic assays of BAR against native and non-native substrates. An apparent KM value of 132±19.2 μM was obtained for phosphinothricin, similar to previously published data^1,3,15^. Aminoadipate and tryptophan are in vitro substrates of BAR but both substrates reached solubility limit before reaching saturation concentration for BAR. Negative controls (open circles) were performed in absence of BAR at the highest substrate concentration tested (20 mM).

To reveal the structural basis for substrate selectivity and catalytic mechanism of BAR that would further enable structure-guided protein engineering, we determined the crystal structures of the BAR/acetyl-CoA holocomplex and the BAR/CoA/phosphinothricin ternary complex (see **Supplementary Table 2** for data collection and refinement statistics). Our refined structures revealed that BAR is an αβ protein harboring a globular tertiary structure resembling the previously reported Gcn5-related N-acetyltransferase (GNAT) structures (**Supplementary Fig. 5**)^16-19^. BAR crystalizes as a homodimer with two active sites symmetrically distributed around the dimer interface inside a large open cavity (**Fig. 4a** and **Supplementary Fig. 6**). The cofactor acetyl-CoA binds to a cleft between α4 and α5 on the opposite side of the dimer interface with the acetyl group pointing toward the catalytic center (**Fig. 4a**). Close examination of the BAR/acetyl-CoA and BAR/CoA/phosphinothricin structures illuminates the catalytic mechanism of BAR (**Fig. 4b, 4c** and **Supplementary Fig. 7**). Similar to other GNATs, BAR utilizes a conserved catalytic Glu88 as a general base to deprotonate the amino group of phosphinothricin through a water molecule as the proton shuttle (**Fig. 4b, 4c**, and **Supplementary Fig. 7**)^19^. The deprotonated amino group then undergoes nucleophilic attack on the carbonyl carbon of acetyl-CoA to produce a tetrahedral intermediate, which is further stabilized by an oxyanion hole composed of a positively charged His137 and its proton donor Tyr107 (**Fig. 4c** and **Supplementary Fig. 7**). It is noteworthy that the structural feature underlying this oxyanion hole in BAR must have arisen independently from the functionally analogous oxyanion hole previously described in the histone acetyltransferase GCN5, featuring a backbone amide nitrogen instead^19^. In the final step of the catalytic cycle, coenzyme A is released from the tetrahedral intermediate as a leaving group to produce acetyl-phosphinothricin (**Fig. 4c**).

**Figure 4.**
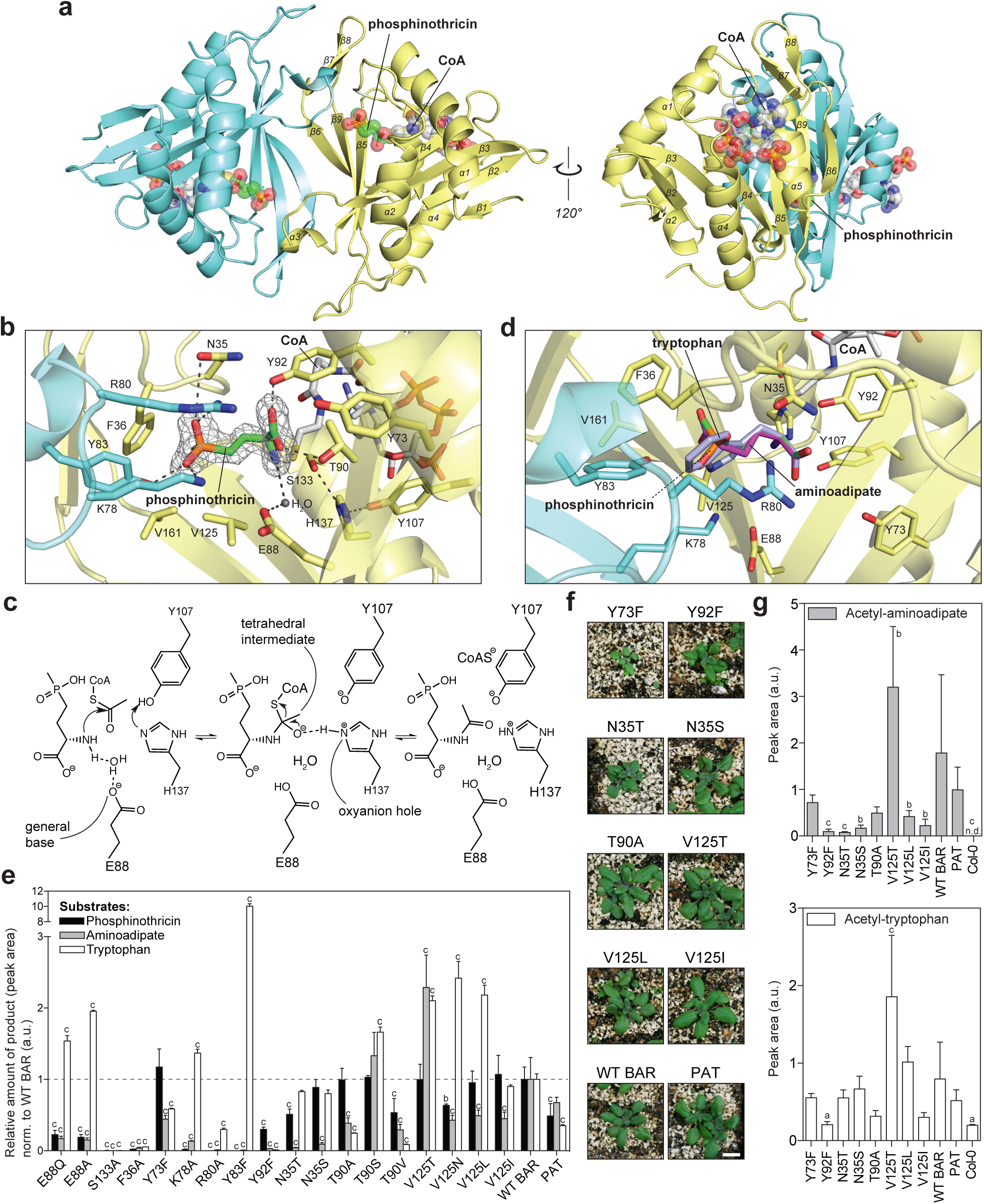
Structural basis for amino acid N-acetylation catalyzed by BAR and structure-guided engineering of BAR with reduced nonspecific activities. (**a**) Cartoon representation of BAR homodimer in complex with phosphinothricin and CoA. Two monomers of the dimer are colored in blue and yellow respectively. (**b**) Close-up view of the BAR active site. The |2Fo-Fc| omit electron density map (contoured at 3.0 σ) is shown for phosphinothricin. (**c**) Proposed catalytic mechanism of BAR. (**d**) Docking of tryptophan and aminoadipate within the BAR active site reveals reduced favorable contacts compared to phosphinothricin. (**e**) Enzyme activity assays using purified BAR mutant proteins against phosphinothricin (0.2mM), aminoadipate (1mM) and tryptophan (1mM). Wild-type BAR (WT BAR) and PAT from *Streptomyces viridochromogenes* were also examined as controls. Assays were terminated during the initial linear rate of product formation. The relative amount of product formed by each BAR mutant was normalized to WT BAR for each substrate (value of 1). Error bars, mean ± s.d. (n = 3 technical replicates). (**f**) Photographs of Arabidopsis T1 lines transformed with select BAR mutants 20 days after phosphinothricin treatment. Scale bar = 0.5 cm. (**g**) Relative levels of acetyl-aminoadipate and acetyl-tryptophan in phosphinothricin-resistant T1 Arabidopsis plants transformed with select BAR mutants. Error bars, mean ± s.d. (n = 4 biological replicates). Significance levels were indicated based on one-way ANOVA with Dunnett's test for multiple comparisons to WT BAR. a, p-value<0.1; b, p-value<0.05; c, p-value<0.01; a.u., arbitrary unit.

The BAR/CoA/phosphinothricin ternary structure also reveals active-site residues involved in phosphinothricin binding. Within each active site, the methylphosphoryl group of the substrate engages hydrophobic interactions with the surrounding Phe36, Gly127, and Val161 from the same monomer, whereas the two phosphoryl oxygen atoms are coordinated by Lys78, Arg80, and Tyr83 from the β3-loop-α3 region of the neighboring monomer via a set of H-bonds and electrostatic interactions (**Fig. 4b**). Furthermore, the amino acid group of phosphinothricin is properly positioned at the catalytic center by an H-bond network involving the backbone carbonyl group of Val125 and the side chains of Thr90 and Tyr92 (**Fig. 4b**). Despite various attempts using co-crystallization and soaking techniques, structures of BAR containing aminoadipate or tryptophan could not be obtained, reflecting the low binding affinity of these nonspecific substrates to BAR. Simulated docking of these substrates within the active site of the BAR/CoA/phosphinothricin structure reveals fewer favorable interactions as well as potential steric clashes with the surrounding residues compared to phosphinothricin (**Fig. 4d**).

Site-directed mutagenesis followed by biochemical assays confirmed the roles of many active-site residues predicted by structural analysis (**Fig. 4e** and **Supplementary Fig. 8**). Mutating the catalytic Glu88 to Ala or Gln greatly reduces the activity of BAR toward phosphinothricin and aminoadipate. Nevertheless, these mutants exhibit higher activity toward tryptophan than that of the wild-type enzyme (**Fig. 4e**), suggesting that tryptophan may be deprotonated through an alternative mechanism independent of Glu88 and/or the first deprotonation step is not rate-limiting for BAR-catalyzed acetyl-tryptophan formation. H137A and Y107A mutants failed to yield soluble recombinant protein (**Supplementary Fig. 8**), preventing the role of the oxyanion hole in catalysis to be directly assessed. We thus probed this indirectly by mutating Ser133, a residue that forms an H-bond with the imidazole ring π-nitrogen of His137 (**Fig. 4b**). The resulting S133A mutant exhibits completely abolished N-acetyltransferase activity toward the three tested substrates, suggesting an essential role of Ser133 in catalysis, likely through proper positioning of the oxyanion hole (**Fig. 4e** and **Supplementary Fig. 7**). Mutants affecting phosphinothricin-binding residues, including F36A, K78A, R80A, Y83F, Y92F, generally show significantly reduced activity toward phosphinothricin and aminoadipate, while K78A and Y83F display increased activity toward the more hydrophobic substrate tryptophan (**Fig. 4e**).

With the structural information of BAR in hand, we sought to engineer BAR through structure-guided mutagenesis to repress its undesired nonspecific activities toward aminoadipate and tryptophan while maintaining its native activity against phosphinothricin. We selected residue positions Asn35, Tyr73, Thr90, Tyr92, and Val125 for targeted mutagenesis based on structural analysis as well as multiple sequence alignment containing BAR, PAT, and other closely related homologs from bacteria (**Fig. 4b, 4d** and **Supplementary Fig. 9**). A set of eleven mutants was first characterized in vitro (**Fig. 4e**), and eight of them were further tested in transgenic Arabidopsis (**Fig. 4f and 4g**). All eight BAR mutants confer phosphinothricin resistance in Arabidopsis (**Fig. 4f** and **Supplementary Fig. 10**). Metabolic profiling of these transgenic lines confirmed that mutations in select active-site residues of BAR can modulate the in vivo nonspecific activities of BAR toward aminoadipate and tryptophan (**Fig. 4g**). Notably, transgenic Arabidopsis plants containing N35T, N35S, V125L or V125I BAR mutants display significantly reduced levels of acetyl-aminoadipate compared to plants containing wild-type BAR (**Fig. 4g**). Moreover, plants expressing Y92F BAR mutant exhibit significantly reduced levels of both acetyl-aminoadipate and acetyl-tryptophan compared to plants containing wild-type BAR. Oppositely, the V125T BAR mutant shows increased promiscuity against both aminoadipate and tryptophan in vitro and in vivo (**Fig. 4e and 4g**). These observed differences in acetyl-aminoadipate and acetyl-tryptophan levels are not due to BAR protein levels in transgenic plants (**Supplementary Fig. 11**), but are consistent with the altered catalytic activities of various BAR mutants measured in vitro (**Fig. 4e** and **Supplementary Fig. 12**). Subsequent kinetic analysis revealed that both N35T and Y92F BAR mutants exhibit compromised affinity toward native substrate phosphinothricin, while retaining largely unaltered catalytic speed compared to wild-type BAR (**Supplementary Fig. 12a**). In contrast, both mutants show drastically reduced catalytic activity toward one or both non-native substrates (**Supplementary Fig. 12b**).

Transgenic expression of enzymes catalyzing a variety of desirable biochemical reactions in heterologous hosts is a common strategy in both basic biological research and translational biotechnology. Prominent examples include reporter enzymes, such as firefly luciferase and β-glucuronidase, antibiotic/herbicide markers, such as aminoglycoside kinase that confers kanamycin resistance and BAR, and many enzymes used for metabolic engineering purposes in microbes and higher eukaryotes^20^. Besides resistance to phosphinothricin, other herbicide-resistance traits employing microbial detoxifying enzymes have also been developed to confer tolerance to alternative herbicides, such as dicamba and bromoxynil^21,22^. Although enzymes are generally considered as perfected catalysts with superior substrate specificity and predictable catalytic mechanism, increasing evidences have raised awareness of the unpredictable behaviors of enzymes and their profound implication in natural and directed evolution of new enzymatic functions^23^. However, whether and how heterologous expression of a foreign enzyme would interfere with the native metabolic system remains an open question to be addressed on a case-by-case basis.

In this study, we discovered that transgenic expression of the herbicide-resistance enzyme BAR of bacterial origin indeed acetylate two endogenous amino acids, resulting in the ectopic accumulation of acetyl-aminoadipate and acetyl-tryptophan. Despite the widespread use of BAR in GM crops^2,24^ and the extensive testing and deregulation processes associated with this trait over the past few decades^1,3,15,25,26^, such phenomenon was not reported earlier, probably due to technological limitation in metabolomics analysis in the past. Our findings therefore suggest a revised procedure incorporating metabolomics analysis for future assessment of GM crops in order to establish substantial equivalence, a central aspect of the risk assessment process^27^. While acetyl-tryptophan is a naturally occurring metabolite found in numerous plant species, including Arabidopsis, *Salsola collina, Glycine max, Solanum lycopersicum, Cocos nucifera*, and *Ginkgo biloba*^28,29^, to the best of our knowledge, acetyl-aminoadipate has never been reported as an endogenous plant metabolite in non-transgenic plants. However, in line with our findings, a recent study showed that acetyl-aminoadipate accumulates in flowers of a BAR-containing mutant of Arabidopsis^30^. Whereas future research is necessary to fully assess the impact of our findings regarding the current use of BAR in GM crops on human health and plant fitness, we provide a solution to reduce the undesirable nonspecific activities of BAR through structure-guided enzyme engineering so that its intended herbicide-degrading activity can be maximally insulated from the metabolome of the host.

## ACKNOWLEDGMENTS

We thank D. M. Sabatini, G. R. Fink, N. Amrhein and E. Martinoia for helpful discussions. We thank J. M. Cheeseman for providing the phosphinothricin-resistant *Glycine max* line. This work is based on research conducted at the Northeastern Collaborative Access Team (NE-CAT) beamlines, which are funded by the National Institute of General Medical Sciences from the National Institutes of Health (P41 GM103403). The Pilatus 6M detector on NE-CAT 24-ID-C beam line is funded by a NIH-ORIP HEI grant (S10 RR029205). This research used resources of the Advanced Photon Source, a U.S. Department of Energy (DOE) Office of Science User Facility operated for the DOE Office of Science by Argonne National Laboratory under Contract No. DE-AC02-06CH11357. This work was supported by the Swiss National Science Foundation (grant 31003A_149389 to S.H. and postdoctoral fellowship P2ZHP3_155258 to B.C.), the EU-funded Plant Fellows program (S.A.), the Pew Scholar Program in the Biomedical Sciences (J.K.W.) and the Searle Scholars Program (J.K.W.).

## AUTHOR CONTRIBUTIONS

B.C., S.A., S.H. and J.K.W. designed experiments; B.C., R.H., L.G., R.F. and S.A. performed experiments; B.C., R.H., L.G. and J.K.W. analyzed data; B.C., S.H., S.A. and J.K.W. wrote the manuscript.

## MATERIALS AND METHODS

### Plant materials

Arabidopsis (*Arabidopsis thaliana*) Columbia-0 (Col-0) and Wassilewskija (Ws) were used as wild types. T-DNA insertion lines were from the following collections: SALK lines^8^: SALK_130606 (SALK_1), SALK_051823C (SALK_2), SALK_110649 (SALK_3); SAIL lines^7^: SAIL_1165_B02 (SAIL_1), SAIL_503_C03 (SAIL_2), SAIL_1235_D10 (SAIL_3); GABI lines^9^: GABI_453E01 (GABI_1), GABI_833F02 (GABI_2), GABI_453A08 (GABI_3); FLAG lines^6^: FLAG_076H05 (*clh2-1*^5^; FLAG_1), FLAG_271B02 (FLAG_2), FLAG_495A09 (FLAG_3), FLAG_271B12 (*FLAG_lkrsdh*). SALK, SAIL and GABI lines were obtained from the European Arabidopsis Stock Center (http://arabidopsis.info/). The FLAG lines were obtained from the INRA Versailles Arabidopsis Stock Center (http://publiclines.versailles.inra.fr/). Homozygous (and heterozygous for FLAG_*lkrsdh*) plants were identified by PCR using T-DNA- and gene-specific primers.

Arabidopsis T-DNA lines used for untargeted metabolomics and relative quantification of acetyl-aminoadipate and acetyl-tryptophan were grown on soil under a 12-h-light/12-h-dark photoperiod with fluorescent light of 80 to 120 μmol photons m^-2^ s^-1^ at 22°C and 60% relative humidity. For senescence induction, leaves from 5-week-old plants were excised and incubated in permanent darkness on wet filter paper for 8 d at ambient temperature. Transgenic Arabidopsis lines transformed with BAR mutants and Arabidopsis T-DNA lines used for absolute quantification of acetyl-aminoadipate and acetyl-tryptophan were grown on soil under a 16-h-light/8-h-dark photoperiod with fluorescent light of 80 to 120 μmol photons m^-2^ s^-1^ at 22°C and 60% relative humidity. For senescence induction, leaves from phosphinothricin-resistant, 4-week-old plants were excised and incubated in permanent darkness on wet filter paper for 6 d at ambient temperature.

Wild-type and phosphinothricin-resistant *Brassica juncea*^31^ was grown on soil under a 16-h-light/8-h-dark photoperiod with fluorescent light of 80 to 120 μmol photons m^-2^ s^-1^ at 22°C and 60% relative humidity. For senescence induction, fully developed cotyledons were excised and incubated in permanent darkness on wet filter paper for 5-7 d at ambient temperature.

Phosphinothricin-resistant *Glycine max* (Liberty Link trait A2704-12, 283 Morril MC-116, Credenz CZ 3841 LL, Bayer CropScience). Wild-type *Glycine max* (variety: Chiba Green) was obtained from local market (High Mowing Organic Seed). *Glycine max* was grown on soil under a 16-h-light/8-h-dark photoperiod with fluorescent light of 80 to 120 μmol photons m^-2^ s^-1^ at 22°C and 60% relative humidity. For testing phosphinothricin resistance, 30-days old plants were sprayed with Finale^®^ (Bayer CropScience) diluted 1:500 in water and photographs were taken after 14 days (**Supplementary Fig. 3**). Green and senescent leaf samples were collected from 40-days old plants.

### Metabolite extraction

Arabidopsis samples were collected in 2 mL Eppendorf tubes containing 500 μL of 1.5 mm glass beads, weighted and snap-frozen in liquid nitrogen. The frozen samples were ground using a MM300 Mixer Mill (Retsch) at 30 Hz for 5 min and stored at -80°C until further processing. *Brassica juncea* and *Glycine max* samples were snap-frozen in liquid nitrogen and ground with a mortar and pestle. Metabolites were extracted using 5 (leaf samples) or 10 to 50 volumes (seed samples; w/v) of ice-cold extraction buffer (80% methanol, 20% water, 0.1% formic acid (v/v/v)). Extracts were homogenized at 30 Hz for 5 min and centrifuged (14,000-16,000 g, 4°C). After re-centrifugation, supernatants were transferred to LC vials and randomly analyzed by LC-MS.

### LC-MS analysis of Arabidopsis T-DNA mutants and *Brassica juncea* (untargeted metabolomics and relative quantification of acetyl-aminoadipate and acetyl-tryptophan)

The LC-MS instrument was composed of an Ultimate 3000 Rapid Separation LC system (Thermo Scientific) coupled to a Bruker Compact ESI-Q-TOF (Bruker Daltonics). The reverse-phase chromatography system consisted of an 150 mm C18 column (ACQUITY UPLC^TM^ BEH, 1.7 μm, 2.1 x 150 mm, Waters), which was developed using LC-MS solvents (Chemie Brunschwig) with a gradient (flow rate of 0.3 mL min^-1^) of solvent B (acetonitrile with 0.1% (v/v) formic acid) in solvent A (water with 0.1% (v/v) formic acid) as follows (all (v/v)): 5% for 0.5 min, 5% to 100% in 11.5 min, 100% for 4 min, 100% to 5% in 1 min and 5% for 1 min. Electrospray ionization (ESI) source conditions were set as follows: gas temperature, 220°C; drying gas, 9 L min^-1^; nebulizer, 2.2 BAR; capillary voltage, 4500 V; end plate offset, 500 V. Tuning conditions were set as follows: funnel 1 RF, 250 Vpp; funnel 2 RF, 150 Vpp; isCID energy, 0 eV; hexapole RF, 50 Vpp; quadrupole ion energy, 3.0 eV; quadrupole low mass, 90 m/z; collision cell, 6 eV; pre-pulse storage time, 3 μs. The instrument was set to acquire over the m/z range 50-1300, with an acquisition rate of 4 spectra s^-1^. Conditions for MS^2^ of automatically selected precursors (data-dependent MS^2^) were set as follows: threshold, 1000 counts; active smart exclusion (5x); active exclusion (exclude after 3 spectra, release after 0.2 min, reconsider precursor if current intensity/previous intensity is ≥5); number of precursors, 3; active stepping (basic mode, timing 50%-50%, collision RF from 350 to 450 Vpp, transfer time from 65 to 80 μs, collision energy from 80 to 120%). All data were recalibrated internally using pre-run injection of sodium formate (10 mM sodium hydroxide in 0.2% formic acid, 49.8% water, 50% isopropanol (v/v/v)). After data recalibration using DataAnalysis (version 4.2, Bruker Daltonics) and data conversion to mzXML format using ProteoWizard MSConvert^32^, metabolite features detected in Ws and FLAG_076H05 (senescent leaves, four replicates) were aligned according to retention time and relatively quantified using XCMS online^33^ (pairwise comparison using XCMS online pre-set parameters “UPLC/Bruker Q-TOF”). Up-regulated features in FLAG_076H05 were identified at retention times of 2.8 min (labeled as “1” in **Fig. 1a**, m/z 204.086 (fold change ≥10, p-value ≤0.005, intensity threshold 800,000)) and 6.5 min (labeled as “2” in **Fig. 1a**, m/z 247.108 (fold change ≥10, p-value ≤0.005, intensity threshold 100,000)) and further characterized as ions derived from N-acetyl-L-aminoadipate and N-acetyl-L-tryptophan, respectively, by database searches in METLIN^34^ using MS and MS^2^ spectra. Relative quantification of acetyl-aminoadipate and acetyl-tryptophan in Arabidopsis mutants from different insertion mutant collections was carried out by QuantAnalysis (version 2.2, Bruker Daltonics) using extracted ion chromatogram (EIC) traces ([M+H]^+^).

### Relative quantification of acetyl-aminoadipate and acetyl-tryptophan in phosphinothricin-resistant *Glycine max*

The LC-MS instrument was composed of an Ultimate 3000 Rapid Separation LC system (Thermo Scientific) coupled to a Q-Exactive mass spectrometer (Thermo Scientific). The reverse-phase chromatography system consisted of an 150 mm C18 Column (Kinetex 2.6 μm silica core shell C18 100Å pore, Phenomenex), which was developed using Optima™ LC/MS solvents (Fisher Chemical) with a gradient (flow rate of 0.8 mL min^-1^) of solvent B (acetonitrile with 0.1% (v/v) formic acid) in solvent A (water with 0.1% (v/v) formic acid) as follows (all (v/v)): 2% for 2 min, 2% to 99% in 25 min, 99% for 5 min, 99% to 2% in 1 min and 2% for 2 min.

The mass spectrometer was configured as to perform 1 MS scan from m/z 100-800 followed by 1-4 data-dependent MS^2^ scans using HCD fragmentation with collision energy of 30 eV. The ion source parameters were as follows: spray voltage (+) at 3000 V, capillary temperature at 275 °C, sheath gas at 40 arb units, aux gas at 15 arb units, sweep gas at 1 arb unit, max spray current at 100 (μA), probe heater temp at 350°C, ion source: HESI-II. Data analysis was performed with Xcalibur (Thermo Scientific) using extracted ion chromatogram (EIC) traces ([M+H]^+^).

### Absolute quantification of acetyl-aminoadipate and acetyl-tryptophan in Arabidopsis T-DNA mutants

Metabolites were extracted as described above and then analyzed on an Ultimate 3000 Rapid Separation LC system (Thermo Scientific) coupled to a TSQ Quantum Access MAX triple-quadrupole mass spectrometer (Thermo Scientific). The reverse-phase chromatography system consisted of an 150 mm C18 column (Kinetex 2.6 μm silica core shell C18 100Å pore, Phenomenex) which was developed using Optima™ LC/MS solvents (Fisher Chemical) with a gradient (flow rate of 0.6 mL min^-1^) of solvent B (acetonitrile with 0.1% (v/v) formic acid) in solvent A (water with 0.1% (v/v) formic acid) as follows (all (v/v)): 2% for 3 min, 2% to 99% in 9 min, 99% for 4 min, 99% to 2% in 1 min and 2% for 1 min. The mass spectrometer was configured to perform two selected-reaction-monitoring scans, each for 0.5 seconds, for acetyl-aminoadipate and acetyl-tryptophan. The m/z resolution of Q1 was set to 0.4 FWHM, the nitrogen collision gas pressure of Q2 was set to 1.5 mTorr, and the Q3 scan width was set to 0.500 m/z in both cases. Selected reaction monitoring for acetyl-aminoadipate was as follows: precursor ion selection at 204.086 m/z on positive ion mode, fragmentation at 10 V, and product ion selection at 144.065 m/z. Selected reaction monitoring for acetyl-tryptophan was as follows: precursor ion selection at 247.107 m/z on positive ion mode, fragmentation at 20 V, and product ion selection at 188.070 m/z. Acetyl-aminoadipate was synthesized using recombinant BAR as described below and used as standard. Pure acetyl-tryptophan was purchased from Sigma-Aldrich.

### Heterologous expression of wild-type BAR and activity determination

The BAR coding sequence was amplified by PCR (KaPa HiFi HotStart polymerase; KaPa Biosystems) from genomic DNA extracted from homozygous plants of the SAIL line SAIL_1165_B02 using primers SAIL_BAR_F_pPROEX and SAIL_BAR_R_pPROEX (see **Supplementary Table 3**) and then cloned into pProEX Hta (Invitrogen) via *Eco*RI and *Hin*dIII resulting in a 6xHis-BAR fusion construct.

6xHis-tagged BAR protein was expressed in *E. coli* BL21(DE3) grown in Terrific Broth medium. At an optical density at 600 nm of 0.6, protein expression was induced with 1.0 mM IPTG and cells were grown at 37°C for 2.5 h. Cells from 1 L culture were harvested by centrifugation and resuspended in 25 mL binding buffer (50 mM Tris-HCl pH 8, 500 mM NaCl, 30 mM imidazole). All the following steps were carried out at 4°C. Cell lysis was performed using a microfluidizer (HC-8000, Microfluidics). The lysate was centrifuged (16,000 g) for 20 min, and the 6xHis-tagged BAR protein was purified by metal affinity (5-ml HisTrap HP column, GE Healthcare) and size-exclusion chromatography (HiLoad 16/600 Superdex 200 pg, GE Healthcare) using an ÄKTA Pure FPLC system (GE Healthcare). The 6xHis-TEV tag was removed from BAR prior to size-exclusion chromatography by overnight incubation with 1 μg of 6xHis-TEV protease^35^ per 10 μg protein in 50 mM Tris-HCl pH 8, 500 mM NaCl, 1 mM dithiothreitol, followed passage through HisTrap HP column. Purified recombinant BAR was dialyzed in storage buffer (12.5 mM Tris-HCl pH 8, 50 mM NaCl, 2 mM dithiothreitol) and concentrated to 13 mg/mL using a ultra-centrifugal filter (10,000 Da MWCO, Amicon EMD Millipore). The purity of recombinant BAR was assessed by SDS-PAGE (**Supplementary Fig. 4a**). Purified BAR was aliquoted, snap-frozen in liquid nitrogen and stored at -80°C until further use.

Enzyme assays were carried out in 2 mM Tris-HCl pH 8 and 10 mM acetyl-CoA (Sigma-Aldrich; final volume 25 μl). Before determining the kinetics of BAR with different substrates, time-dependent activity of the purified protein was tested at substrate concentrations of 500 μM L-phosphinothricin (glufosinate ammonium, considered as a 1:1 mixture of L-and D-enantiomers; Sigma-Aldrich) or 1 mM (L-aminoadipate and L-tryptophan; Sigma-Aldrich). Reactions were initiated by the addition of purified BAR at 0.26 μM (assays with L-phosphinothricin) or 150 μM (assays with aminoadipate or tryptophan) and incubated at 25°C for the indicated times (**Supplementary Fig. 4b-d**). Reactions were stopped by the addition of four volumes of 10% water, 90% acetonitrile (v/v), 5 mM ammonium formate pH 3. Likewise, substrate concentration-dependence was determined by incubating assays for 25 min (assays with L-phosphinothricin), 3 h (assays with aminoadipate) or 7 h (assays with tryptophan; **Fig. 3**). Stock solutions of aminoadipate and tryptophan at 60mM were made in 2 mM Tris-HCl pH 8 supplemented with 1 mM N-nonyl β-D-glucopyranoside and substrate concentration-dependence assays employing these two substrates contained 0.33 mM N-nonyl β-D-glucopyranoside. Control assays (**Fig. 3**) were performed with aminoadipate and tryptophan at 20 mM, but in the absence of BAR.

The assays were analyzed on an Ultimate 3000 Rapid Separation LC system (Thermo Scientific) coupled to a TSQ Quantum Access MAX triple-quadrupole mass spectrometer (Thermo Scientific). Assays on phosphinothricin were analyzed as follows. The normal-phase chromatography system consisted of an 150 mm HILIC column (Kinetex 2.6 μm silica core shell HILIC 100Å pore, Phenomenex), which was developed using Optima™ LC/MS solvents (Fisher Chemical) with a gradient (flow rate of 0.8 mL min^-1^) of solvent B (50% water, 50% acetonitrile (v/v), 5 mM ammonium formate pH 3) in solvent A (10% water, 90% acetonitrile (v/v), 5 mM ammonium formate pH 3) as follows (all (v/v)): 0% for 2 min, 0% to 70% in 10 min, 70% to 100% in 30 sec, 100% for 90 sec, 100% to 0% in 30 sec and 0% for 3.5 min. The mass spectrometer was configured to perform selected-ion-monitoring scans of 0.5 seconds using Q3 (center mass m/z: 224.068, scan width 1.0 m/z, scan time 0.5 sec). Assays on aminoadipate and tryptophan were analyzed as described above for the absolute quantification of acetyl-aminoadipate and acetyl-tryptophan in planta. Product formation was quantified using standards synthesized using recombinant BAR (acetyl-phosphinothricin and acetyl-aminoadipate) or commercially available (acetyl-tryptophan, Sigma-Aldrich). *K_M_* and *V*max value for phosphinothricin were inferred using the Michaelis-Menten kinetics nonlinear regression function under Prism 6 (GraphPad).

### X-ray crystallography

Purified BAR protein was incubated with 1 mM acetyl-CoA for >2 hour prior to setting crystal trays. Crystals of BAR were obtained after 3 days at 20 °C in hanging drops containing 1 μL of protein solution (7.5 mg/mL) and 1 μL of reservoir solution (0.18 M calcium acetate, 0.1 M Tris-HCl pH 7, 18% (w/v) PEG 3000, 0.2% (v/v) N-nonyl β-D-glucopyranoside, 1 mM acetyl-CoA). Several crystals were soaked in reservoir solution supplemented with 30 mM L-phosphinothricin for 30-60 min before freezing. Crystals were frozen in reservoir solution supplemented with 15% (v/v) ethylene glycol. Acetylation of phosphinothricin occurred during soaking as no density for the acetyl group of acetyl-CoA was observed in the BAR/CoA/phosphinothricin ternary complex. Co-crystallization screens and soaking experiments were performed using aminoadipate and tryptophan at final concentrations between 5-30 mM. However, we could not identify electron density supporting the presence of these substrates within the resolved crystal structures.

X-ray diffraction data were collected on the 24-ID-C beam line of the Structural Biology Center at the Advanced Photon Source (Argonne National Laboratory) equipped with a Pixel Array Detector (Pilatus-6MF). Diffraction intensities were indexed, integrated, and scaled with the iMosflm^36^ and SCALA^37^ programs. Initial phases were determined by molecular replacement using Phaser under Phenix^38^. Subsequent structural building and refinements utilized Phenix programs^38^. Coot was used for graphical map inspection and manual rebuilding of atomic models^39^. Crystallographic calculations were performed using Phenix. Molecular graphics were produced with the program PyMol.

### Heterologous expression of BAR mutants and activity determination

Single amino acid mutants of BAR were generated using the QuikChange II site-directed mutagenesis kit (Agilent Technologies) and 6xHis-BAR in pProEX Hta as template (see **Supplementary Table 3** for primer sequences). PAT from *Streptomyces viridochromogenes* was amplified using primers BAC0327 and BAC0328 from pAG31 vector^40^ (Addgene 35124) and cloned into *Bam*HI/*Hin*dIII-linearized pProEX Hta by Gibson assembly (New England Biolabs). Wild-type 6xHis-BAR, 6xHis-BAR mutants and 6xHis-PAT were expressed in *E. coli* BL21(DE3) grown in Terrific Broth medium. At an optical density at 600 nm of 0.6, protein expression was induced with 1.0 mM IPTG and cells were grown at 37°C for 2.5 h. Cells from a 150 mL cultures were harvested by centrifugation, lysed using B-PER™ Bacterial Protein Extraction Reagent (Thermo Scientific) and purified by metal affinity using Ni-NTA Agarose (Qiagen). Purified recombinant proteins were concentrated and buffer-exchanged using storage buffer (10 mM Tris-HCl pH 8.0, 0.2 M NaCl, 10% (v/v) glycerol, 1 mM dithiothreitol) and ultra-centrifugal filters (10,000 Da MWCO, Amicon EMD Millipore). The purity of the recombinant proteins was assessed by SDS-PAGE. Final protein concentrations were determined and normalized using a NanoDrop 2000 UV-VIS spectrometer (Thermo Scientific).

Enzyme assays for comparing the relative activity of the purified BAR mutants were carried out in 2 mM Tris-HCl pH 8 and 5 mM acetyl-CoA (Sigma-Aldrich) (final reaction volume 12 μL). Reactions were initiated by the addition of purified recombinant protein at 0.2 μM (assays with L-phosphinothricin at 0.2 mM) or 150 μM (assays with aminoadipate or tryptophan at 1 mM) and incubated at 25°C for 15 min (phosphinothricin), 165 min (aminoadipate), or 330 min (L-tryptophan). Substrate concentration-dependences toward phosphinothricin, aminoadipate and tryptophan were determined for the BAR mutants Y92F and N35T in 2 mM Tris-HCl pH 8 and 10 mM acetyl-CoA (Sigma-Aldrich). Note that assays on aminoadipate and tryptophan were supplemented with 0.33 mM of N-nonyl β-D-glucopyranoside (see also above). Reactions were stopped by the addition of four volumes of 10% water, 90% acetonitrile (v/v), 5 mM ammonium formate pH 3, centrifuged for 2 min (14,000-16,000 g), and transferred to LC vials.

The assays were analyzed on an Ultimate 3000 Rapid Separation LC system (Thermo Scientific) coupled to a TSQ Quantum Access MAX triple-quadrupole mass spectrometer (Thermo Scientific). Assays on phosphinothricin were analyzed as described above. Assays on aminoadipate were analyzed as follows. The reverse-phase chromatography system consisted of an 150 mm C18 column (Kinetex 2.6 μm silica core shell C18 100Å pore, Phenomenex), which was developed using Optima™ LC/MS solvents (Fisher Chemical) with a gradient (flow rate of 0.6 mL min^-1^) of solvent B (acetonitrile with 0.1% (v/v) formic acid) in solvent A (water with 0.1% (v/v) formic acid) as follows (all v/v): 1% for 2 min, 1% to 30% in 9 min, 30% to 99% in 30 sec, 99% for 30 sec, 99% to 1% in 1 min and 1% for 2 min. The mass spectrometer was configured to perform selected-ion-monitoring scans of 0.5 seconds using Q3 (center mass m/z: 204.086, scan width 0.5 m/z, scan time 0.5 sec). Assays on tryptophan were analyzed as follow: the reverse-phase chromatography system consisted of an 150 mm C18 column (Kinetex 2.6 μm silica core shell C18 100Å pore, Phenomenex) which was developed using Optima™ LC/MS solvents (Fisher Chemical) with a gradient (flow rate of 0.7 mL min^-1^) of solvent B (acetonitrile with 0.1% (v/v) formic acid) in solvent A (water with 0.1% (v/v) formic acid) as follows (all v/v): 5% for 1 min, 5% to 99% in 9 min, 99% for 2 min, 99% to 5% in 2 min and 5% for 1 min. The mass spectrometer was configured to perform selected-ion-monitoring scans of 0.5 seconds using Q3 (center mass m/z: 247.108, scan width 0.5 m/z, scan time 0.5 sec).

### Analysis of BAR mutants in planta

Wild-type BAR from *Streptomyces hygroscopicus*, selected BAR mutants and wild-type PAT from *Streptomyces viridochromogenes* were amplified by PCR (Phusion polymerase; New England Biolabs) from pProEX Hta clones (see above) using primers listed in **Supplementary Table 3** and cloned into *Bpi*I-linearized pICH41308^41^ (Golden Gate entry vector) by Gibson assembly (New England Biolabs). BAR and PAT coding sequences were fused with *Agrobacterium tumefaciens* mannopine synthase promoter (from pICH85281) and terminator (from pICH77901) into the empty binary vector pICH47732 by Golden Gate assembly^41^. pICH47732 constructs were transformed into *Agrobacterium tumefaciens* GV3130 strain by electroporation and transformed into Arabidopsis Col-0 by the floral dip method^42^. T1 plants were grown on soil and transformants were selected with Finale^®^ (contains 11.33% glufosinate ammonium; Bayer CropScience) diluted 1:500 in water. Photographs were taken 20 days after herbicide treatment (**Supplementary Fig. 10**).

Metabolites were extracted from dark-incubated leaves collected from T1 phosphinothricin-resistant individuals and then analyzed as described above for the absolute quantification of acetyl-aminoadipate and acetyl-tryptophan in Arabidopsis T-DNA mutants and *Glycine max*.

Protein levels of the BAR mutants in T2 lines were measured as follow. For each protein extraction, equal amounts of aerial tissues from 5-6 T2 populations grown from seeds from independent T1 plants were pooled. Total proteins were isolated from frozen samples by homogenization in 5 volumes of ice-cold extraction buffer [50 mM Tris–HCl pH 8, 100 mM NaCl, 0.5% (v/v) TritonX-100, 2mM β-mercaptoethanol] complemented with a protease inhibitor cocktail (Complete; Roche Diagnostics). Samples were centrifuged at 12,000 g for 5 min and protein concentration of the supernatant was determined using the Bradford Assay (Bio-Rad). Proteins were subsequently precipitated with chloroform–methanol and 10 μg were analyzed by SDS-PAGE and immunoblotting as described^43^. The following antibodies were used for immunoblot analysis: a primary polyclonal antibody against BAR from *Streptomyces hygroscopicus* produced in rabbit (1:1000; P0374-Sigma-Aldrich) and a polyclonal horseradish peroxidase conjugated goat anti-rabbit IgG as the secondary antibody (1:50000; A0545-Sigma-Aldrich). Substrate detection was performed by chemiluminescence (ECL Western Blotting Substrate™ (Pierce)) and *film* exposure.

## SUPPLEMENTARY INFORMATIONS

**Supplementary Table 1.** Absolute quantification of acetyl-aminoadipate and acetyl-tryptophan in senescent leaves and seeds of Arabidopsis

| **Arabidopsis – Senescent leaves** |  |  |
| --- | --- | --- |
|  | Acetyl-aminoadipate (µg/g FW) | Acetyl-tryptophan (µg/g FW) |
| **Col-0** | n.d. | 0.20 ± 0.0 |
| **SAIL-1** | 85.2 ± 7.7 | 4.11 ± 0.6 |
| **SAIL-2** | 80.4 ± 9.9 | 3.75 ± 0.2 |
| **SAIL-3** | 71.2 ± 9.5 | 3.28 ± 0.7 |
| **Ws** | n.d. | 0.20 ± 0.0 |
| **FLAG-1** | 232.8 ± 24.6 | 23.96 ± 1.1 |
| **FLAG-2** | 198.2 ± 15.3 | 21.32 ± 2.3 |
| **FLAG-3** | 307.7 ± 36.3 | 27.83 ± 3.3 |

| **Arabidopsis – Seeds** |  |  |
| --- | --- | --- |
|  | Acetyl-aminoadipate (µg/g FW) | Acetyl-tryptophan (µg/g FW) |
| **Col-0** | n.d. | 0.14 ± 0.06 |
| **SAIL-1** | 42.7 ± 11.7 | 0.34 ± 0.11 |
| **SAIL-2** | 43.4 ± 12.0 | 0.26 ± 0.09 |
| **SAIL-3** | 59.8 ± 1.9 | 0.35 ± 0.04 |
| **Ws** | n.d. | 0.13 ± 0.08 |
| **FLAG-1** | 117.4 ± 48.0 | 3.06 ± 0.47 |
| **FLAG-2** | 237.8 ± 29.9 | 3.40 ± 0.33 |
| **FLAG-3** | 125.8 ± 10.6 | 1.18 ± 0.30 |

Mean ± s.d. (n = 3 biological replicates); FW, fresh weight; n.d., not detected

**Supplementary Table 2.** Data collection and refinement statistics

Data collection and refinement statistics
|  | BAR-ACO (PDB ID: 5T7D) | BAR-COA-Phosphinothricin (PDB ID: 5T7E) |
| --- | --- | --- |
| Resolution range (Å) | 44.79 - 1.4 (1.45 - 1.4) | 62.52 - 1.8 (1.864 - 1.8) |
| Space group | P 1 21 1 | P 1 21 1 |
| Unit cell (Å)<br>(°) | 65.1 71.5 84.05 90 104.33 90 | 64.18 66.11 86.15 90 103.06 90 |
| Total reflections | 238978 (13056) | 93996 (4082) |
| Unique reflections | 131315 (10035) | 54563 (3385) |
| Multiplicity | 1.8 (1.3) | 1.7 (1.2) |
| Completeness (%) | 89.48 (68.67) | 83.62 (52.15) |
| Mean I/sigma(I) | 8.84 (0.98) | 5.41 (1.03) |
| Wilson B-factor (Å<sup>2</sup>) | 14.16 | 16.54 |
| R-merge | 0.03773 (0.5096) | 0.07711 (0.3945) |
| R-meas | 0.05336 | 0.109 |
| CC1/2 | 0.998 (0.611) | 0.96 (0.781) |
| CC* | 1 (0.871) | 0.99 (0.936) |
| R-work | 0.1611 (0.3214) | 0.2094 (0.3383) |
| R-free | 0.1931 (0.3451) | 0.2421 (0.3723) |
| Number of non-hydrogen atoms | 6939 | 6246 |
| macromolecules | 5659 | 5551 |
| ligands | 236 | 278 |
| water | 1044 | 417 |
| Protein residues | 706 | 708 |
| RMS(bonds) (Å) | 0.007 | 0.003 |
| RMS(angles) (°) | 1.27 | 0.91 |
| Ramachandran favored (%) | 99 | 98 |
| Ramachandran outliers (%) | 0 | 0.14 |
| Clashscore | 5.91 | 4.81 |
| Average B-factor (Å<sup>2</sup>) | 20.9 | 22.4 |
| macromolecules | 18.4 | 21.5 |
| ligands | 21 | 28.1 |
| solvent | 34.4 | 30.3 |

Values in parentheses are for highest resolution shell.

**Supplementary Table 3.**
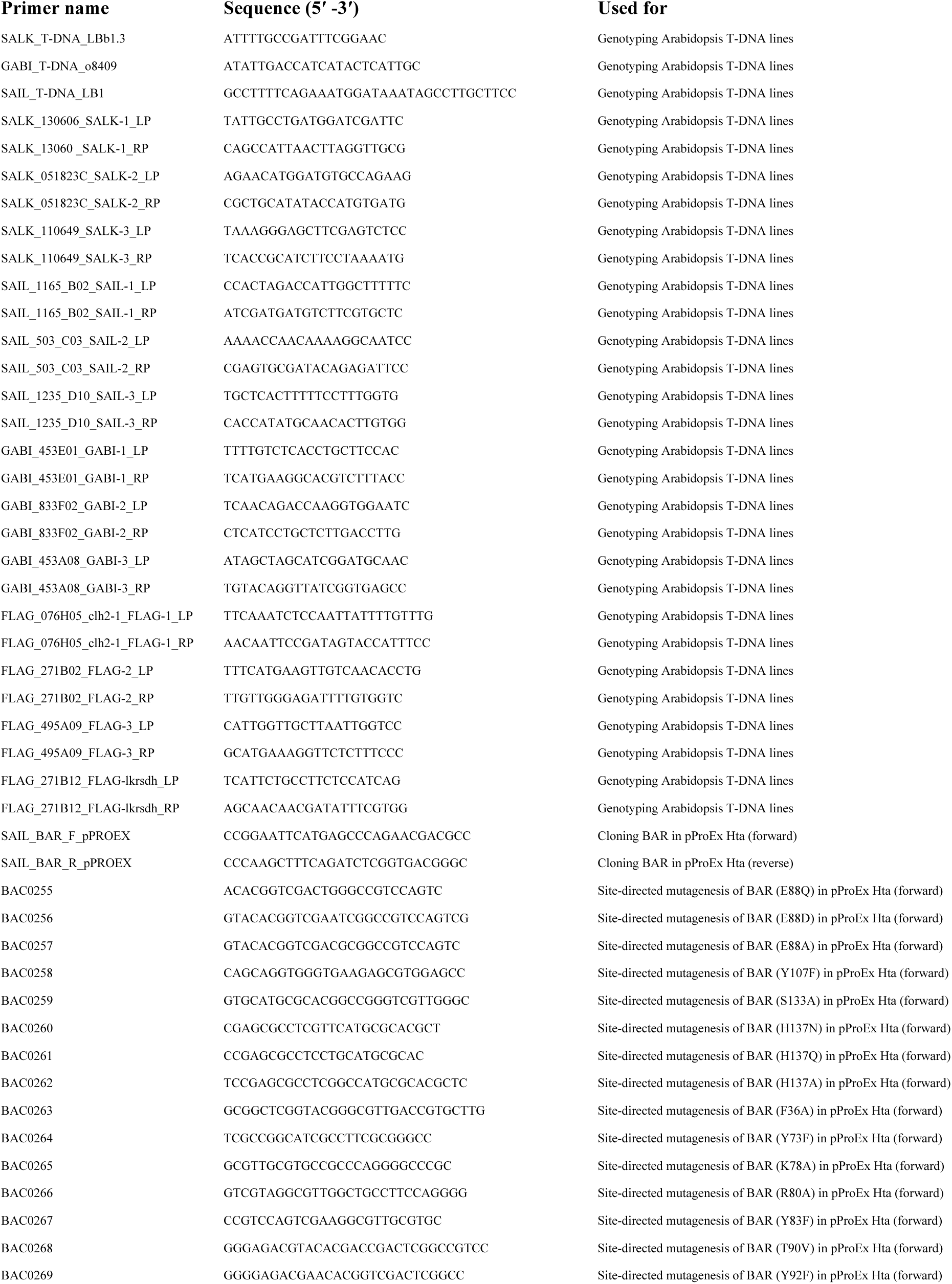
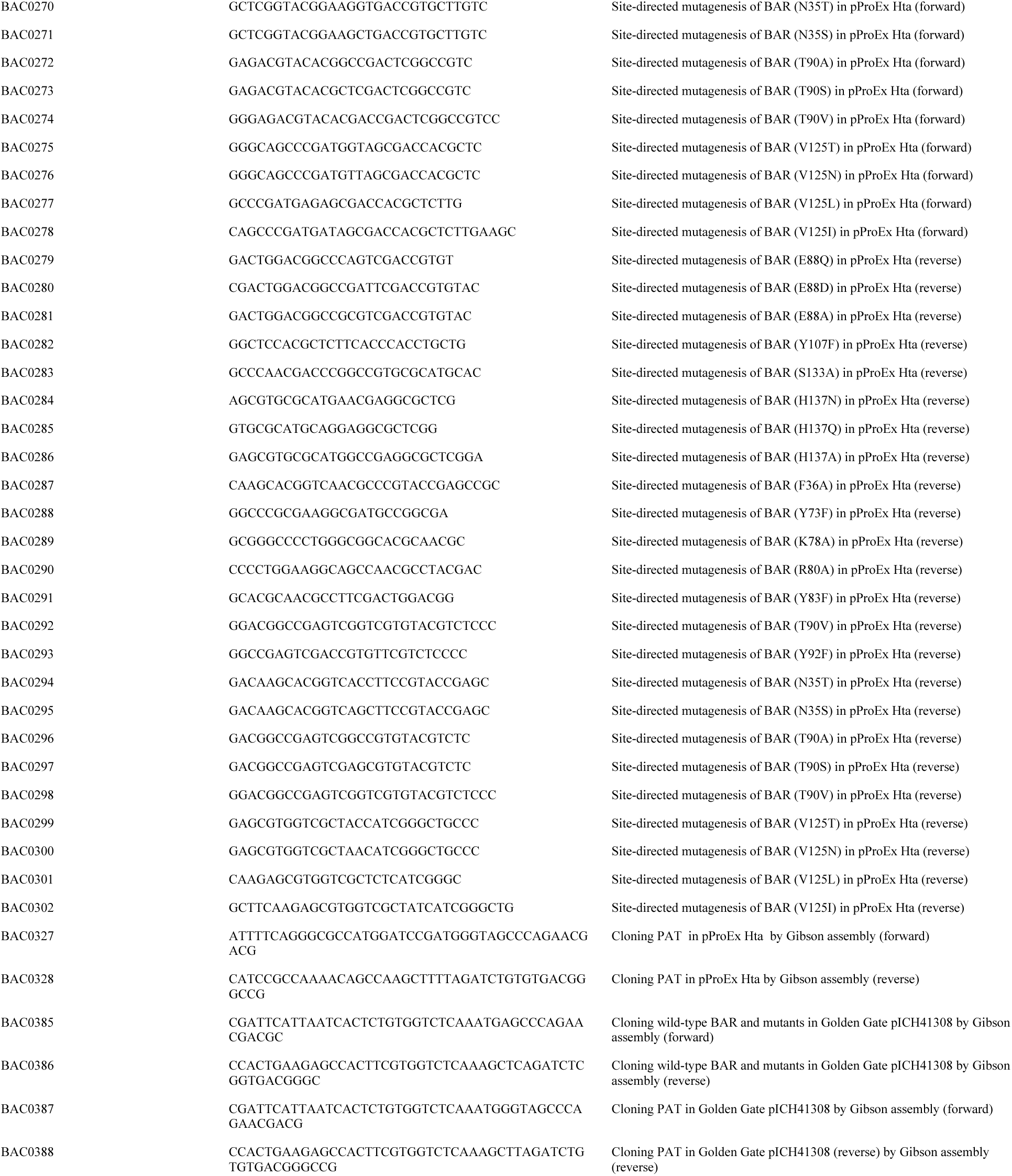
List of primers used in this study

**Supplementary Figure 1.**
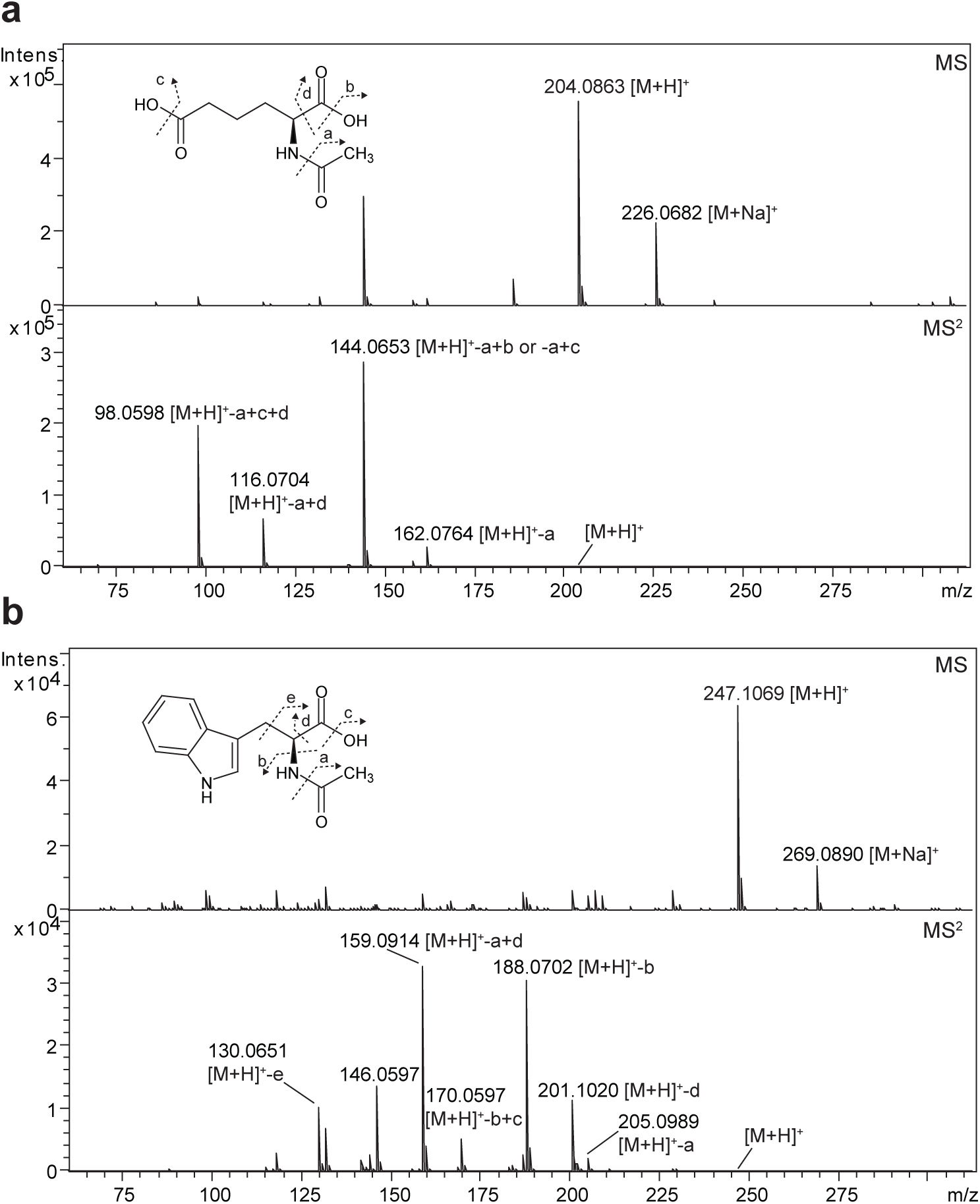
Liquid chromatography-tandem mass spectrometry (LC-MS2) identification of N-acetyl-L-aminoadipate (**a**) and N-acetyl-L-tryptophan (**b**) accumulating in senescent leaves *clh2-1* (FLAG_076H05). MS and MS^2^ spectra, chemical structures and fragmentation pattern are shown. Note that the fragmentation pattern for N-acetyl-L-aminoadipate and N-acetyl-L-tryptophan is concordant with the fragmentation pattern of L-aminoadipate, L-tryptophan and N-Acetyl-DL-tryptophan as in METLIN database (https://metlin.scripps.edu/index.php^34^) and as published previously^29^.

**Supplementary Figure 2.**
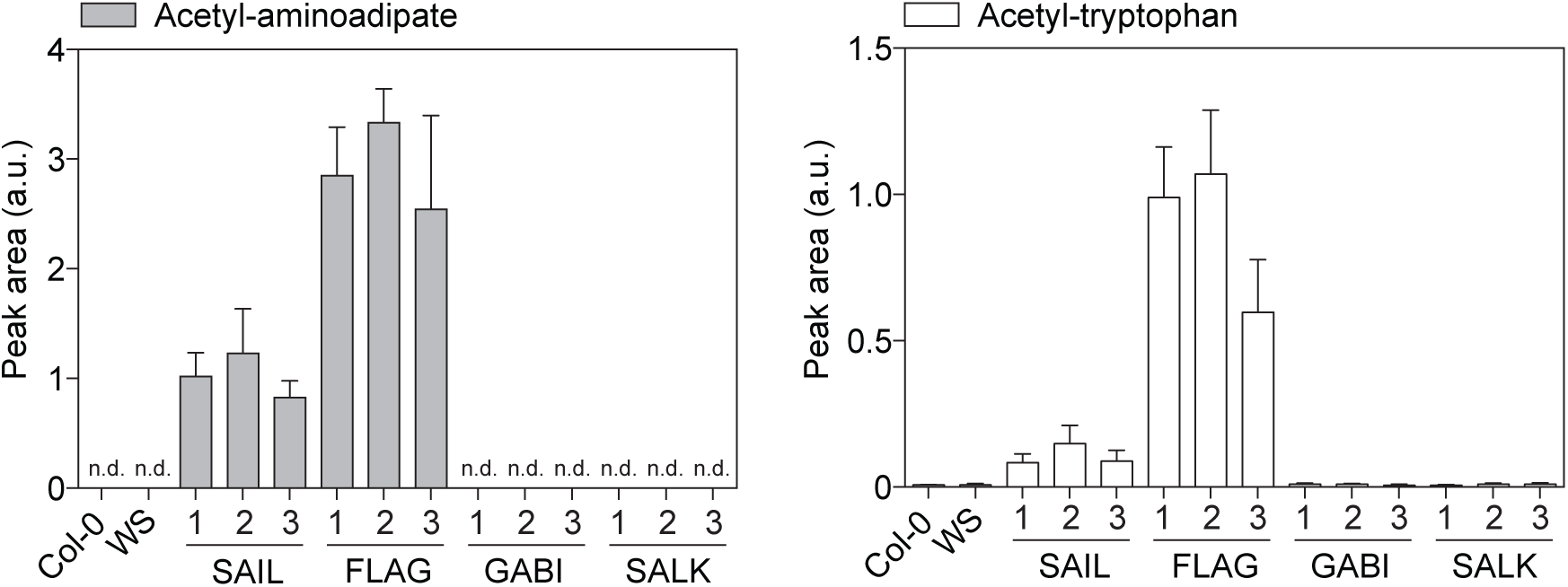
Relative quantification of acetyl-aminoadipate and acetyl-tryptophan in seeds of Arabidopsis mutants from different insertional mutant collections that contains either the *BAR* gene (SAIL and FLAG) or alternative selection marker genes (SALK and GABI). Error bars, mean ± s.d. (n = 6 biological replicates). See **Supplementary Table 1** for absolute quantification. a.u., arbitrary unit; n.d., not detected.

**Supplementary Figure 3.**
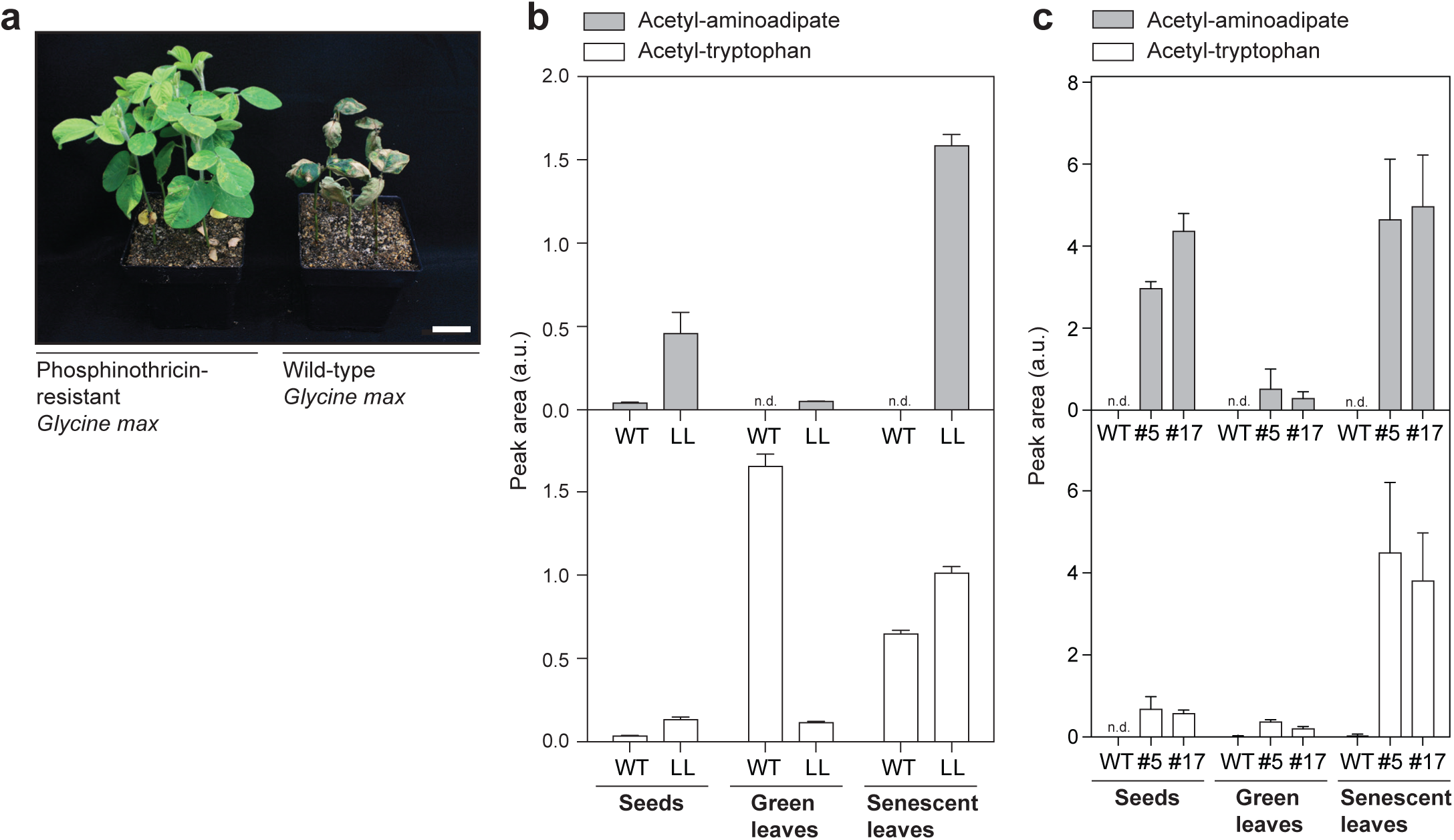
Accumulation of acetyl-aminoadipate and acetyl-tryptophan in *Glycine max* and *Brassica juncea* lines that are resistant to phosphinothricin. (**a**) Photograph of wild-type and phosphinothricin-resistant *Glycine max* taken 14 days after herbicide treatment. Scale bar = 2 cm. (**b**) Relative quantification of acetyl-aminoadipate and acetyl-tryptophan in seeds, green leaves and senescent leaves of wild-type and phosphinothricin-resistant *Glycine max* (LL). Note that wild-type *Glycine max* has been shown to accumulate acetyl-tryptophan naturally^29^. (**c**) Relative quantification of acetyl-aminoadipate and acetyl-tryptophan in seeds, green leaves and senescent leaves of wild-type and phosphinothricin-resistant *Brassica juncea* (lines #5 and #17^31^). Error bars, mean ± s.d. (n = 3 biological replicates). a.u., arbitrary unit.

**Supplementary Figure 4.**
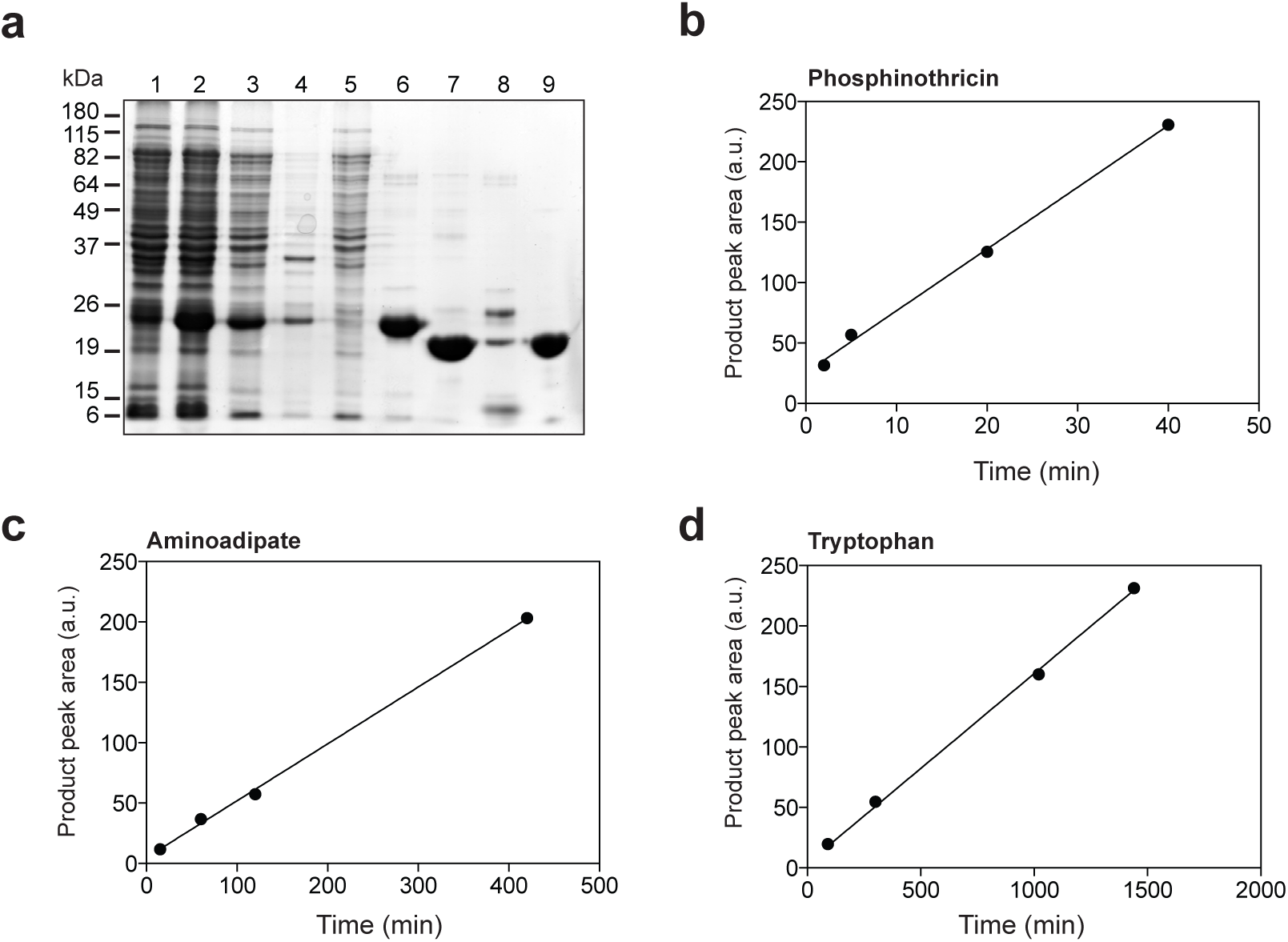
Purification and time-dependent activities of recombinant BAR from *E.coli*. (**a**) BAR expression and purification was monitored by SDS-PAGE. The 6xHis-BAR protein fusion was isolated from the *E. coli* lysate (lane 1, uninduced cells; lane 2, induced cells; lane 3, soluble proteins; lane 4, insoluble proteins) by metal affinity chromatography (Ni^2+^-charged HisTrap (GE Healthcare)); lane 5, flow-through; lane 6, 6xHis-BAR elution). Partially purified 6xHis-BAR protein fusion was then treated with 6xHis-TEV protease^35^ and passed through the HisTrap to remove the His-tag (lane 7, flow-through; lane 8, elution of 6xHis-TEV and uncut 6xHis-BAR) and further purified by gel exclusion chromatography (lane 9). Time-dependent activities of purified 6xHis-BAR were determined at substrate concentration of 500 μΜ for phosphinothricin (**b**) and 1000 μΜ for aminoadipate (**c**) and tryptophan (**d**). a.u., arbitrary unit.

**Supplementary Figure 5.**
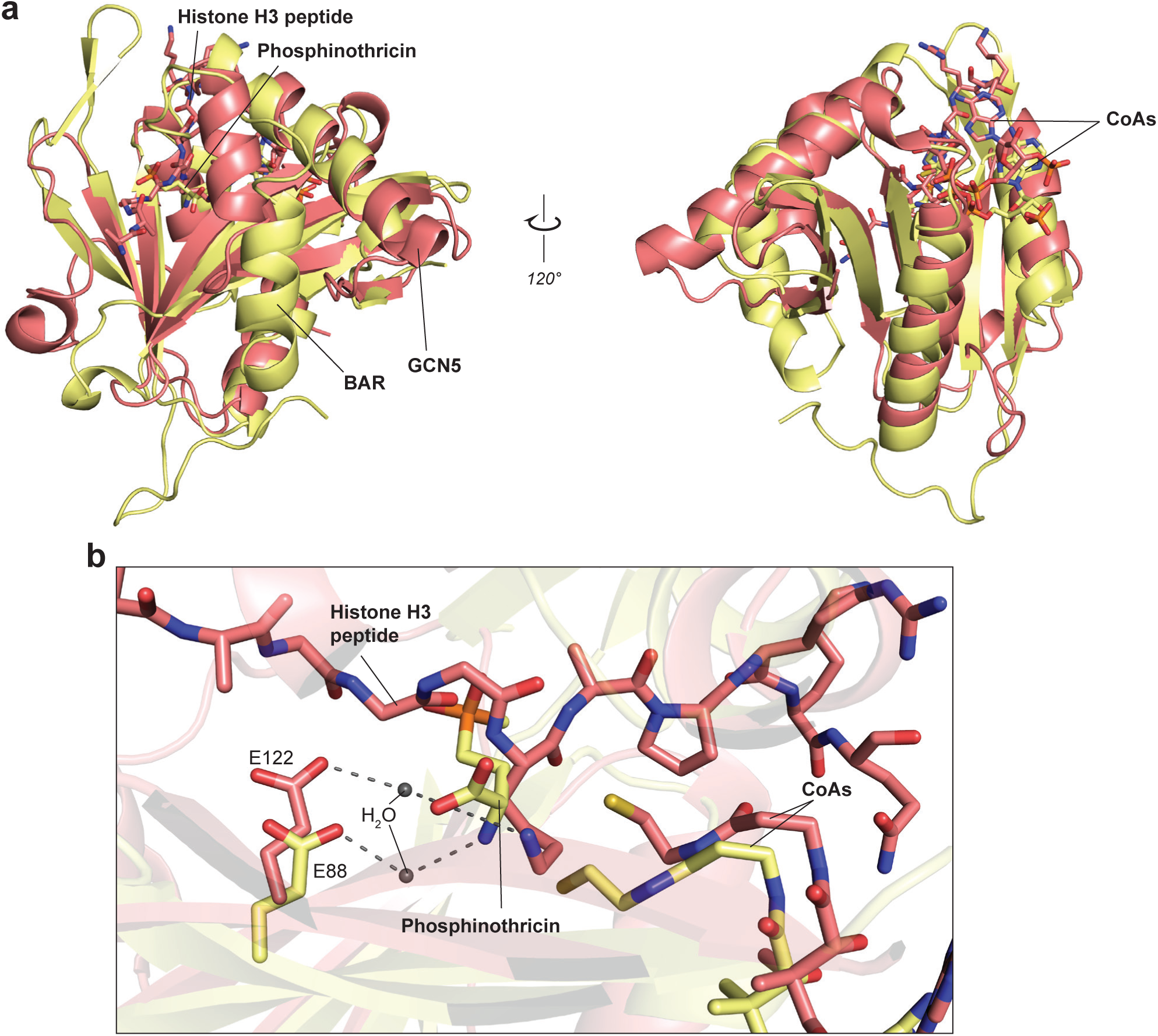
Structural alignment of BAR/CoA/phosphinothricin ternary complex (yellow) and *Tetrahymena* GCN5 bound to CoA and histone H3 peptide (red, PBD ID: 1QSN). (**a**) Diagram showing two views of the alignment perfomed using the SSM structural alignment function under Coot^39^. (**b**) Close-up view of the active site.

**Supplementary Figure 6.**
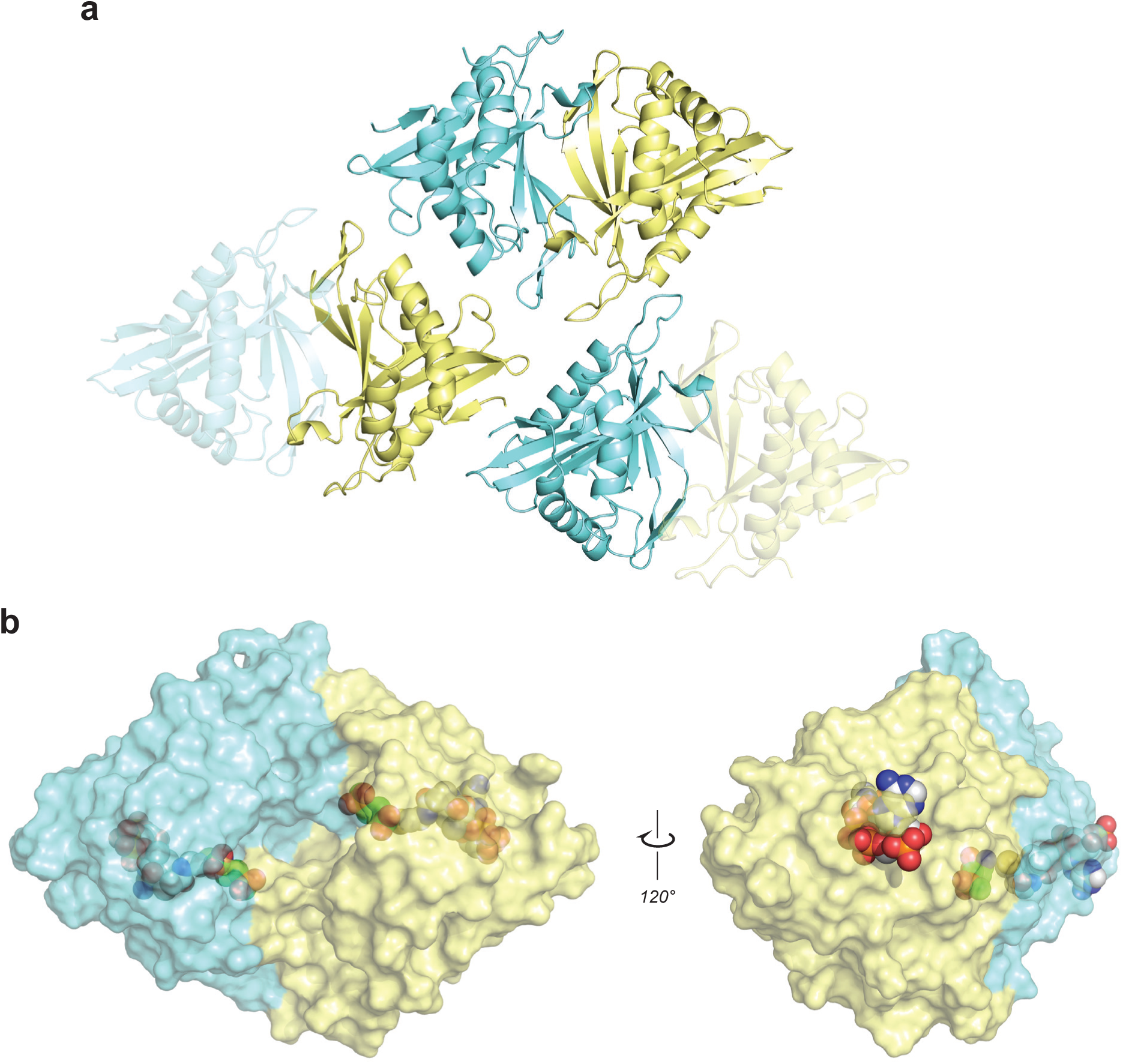
BAR crystallizes as homodimer with two active sites symmetrically distributed around the dimer interface. (**a**) Each asymmetric unit (ASU) is constituted of one homodimer and two monomers that form homodimer with chains from neighboring cells (shown as transparent chains). (**b**) Surface representation of BAR revealing a large open cavity at the dimer interface.

**Supplementary Figure 7.**
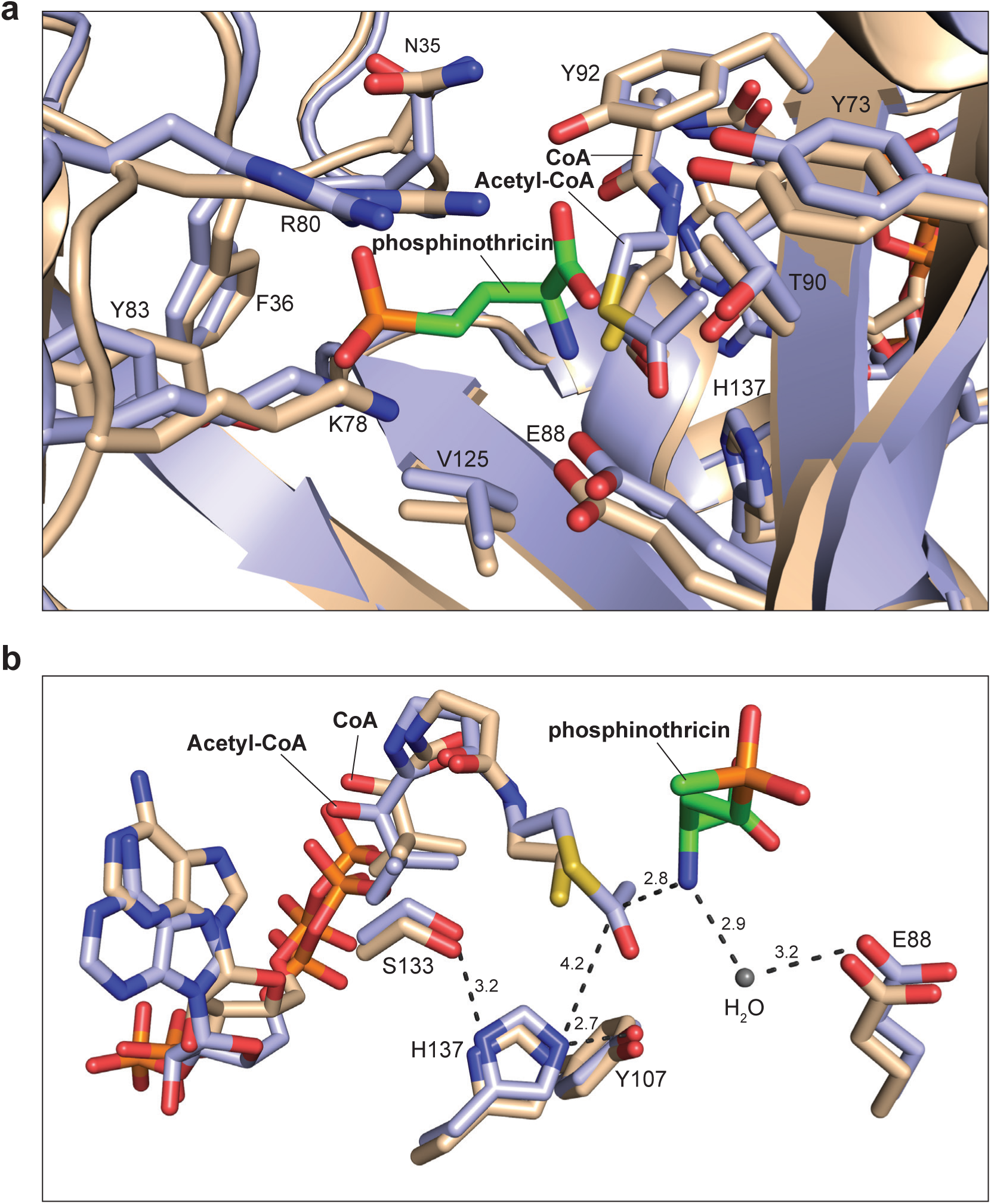
Structural alignment of the BAR/acetyl-CoA holocomplex (purple) with the BAR/CoA/phosphinothricin ternary complex (brown). (**a**) Close-up of view of the active site of BAR. (**b**) Diagram showing the residues involved in catalysis. Distances are shown in Angstroms.

**Supplementary Figure 8.**
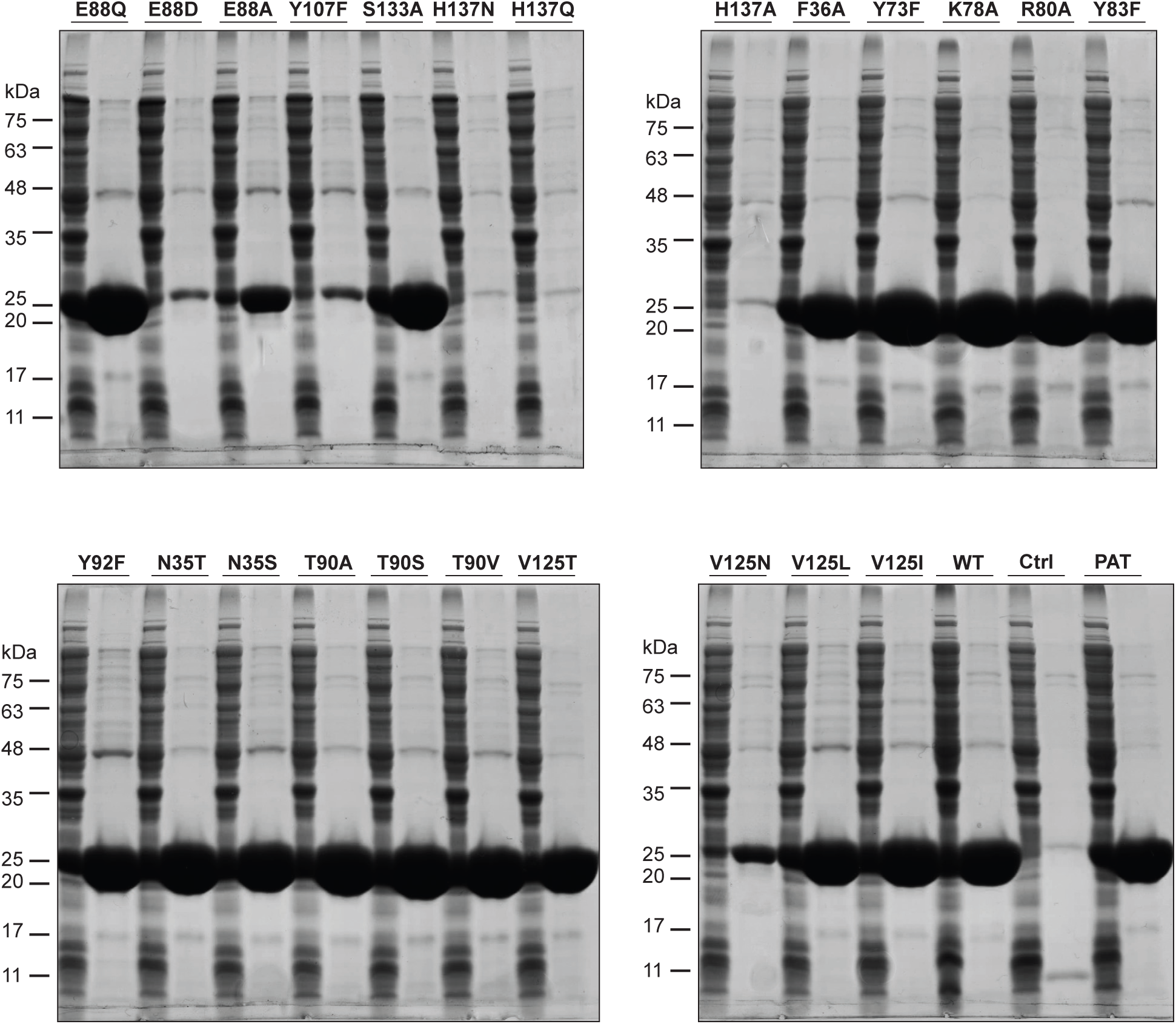
Expression and purification of 23 recombinant mutant versions of BAR from *Streptomyces hygroscopicus* and wild-type PAT from *Streptomyces viridochromogenes*. Left SDS-PAGE lane, soluble fraction of *E.coli* lysate; right lane, purified protein; Ctrl, empty vector.

**Supplementary Figure 9.**
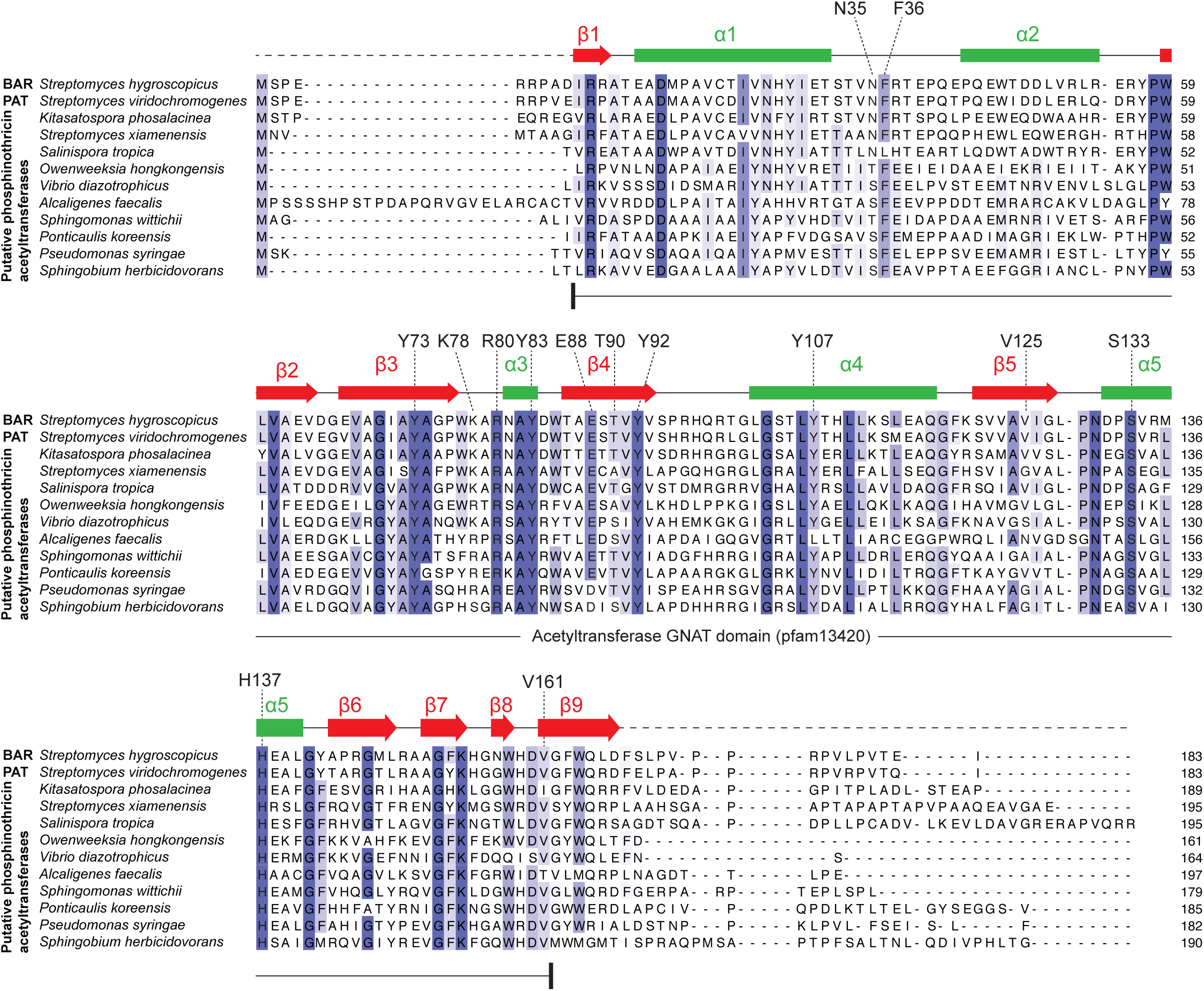
Protein sequence alignement of BAR from *Streptomyces hygroscopicus*, PAT from *Streptomyces viridochromogenes* and closely related homologues from other species. Active site residues as displayed in **Fig. 4b** are labelled. The alignment was performed using Jalview V2 (T-Coffee, default settings^44^). Secondary structure of BAR as labelled in **Fig. 4a** is shown. The acetyltransferase GNAT domain (pfam13420) is displayed. Protein sequences related to BAR from *Streptomyces hygroscopicus* were retrieved from GenBank at the NCBI website using protein BLAST search (http://blast.ncbi.nlm.nih.gov/Blast.cgi). Protein sequence accessions (GenBank): *Streptomyces hygroscopicus*: CAA29262; *Streptomyces viridochromogenes*, WP_003988626; *Kitasatospora phosalacinea*, WP_033213694; *Streptomyces xiamenensis*, AKG45686; *Salinispora tropica*, WP_028566484; *Owenweeksia hongkongensis*, WP_014202881; *Vibrio diazotrophicus*, WP_042485812; *Alcaligenes faecalis*, CAA00175; *Sphingomonas wittichii*, WP_037526498; *Ponticaulis koreensis*, WP_022694195; *Pseudomonas syringae*, WP_032656505; *Sphingobium herbicidovorans*, WP_037462269.

**Supplementary Figure 10.**
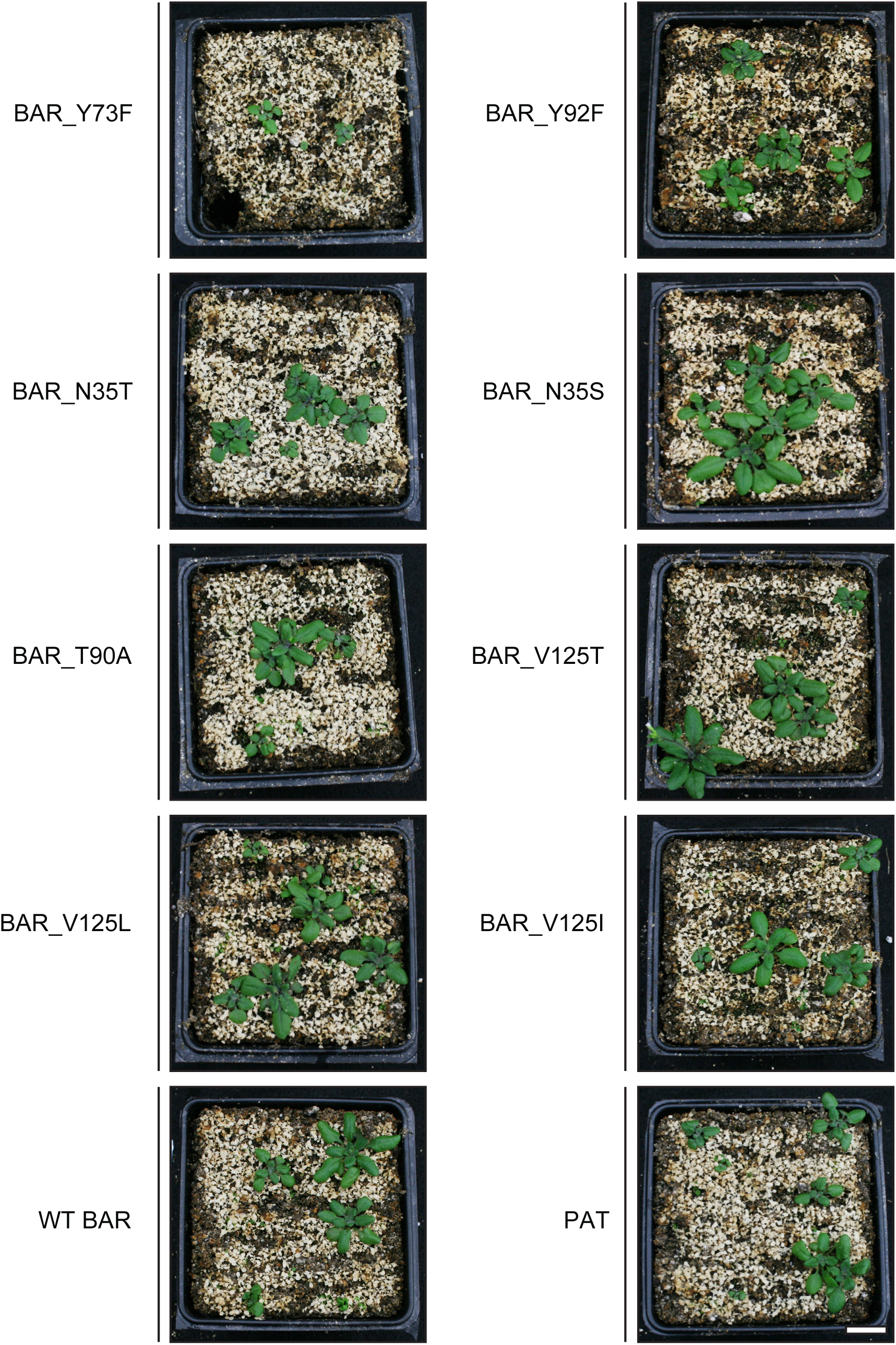
Selected BAR variants confer resistance to phosphinothricin in Arabidopsis. Photographs of Arabidopsis T1 lines transformed with wild-type BAR from *Streptomyces hygroscopicus* (WT BAR), PAT from *Streptomyces viridochromogenes* and selected BAR mutants taken 20 days after phosphinothricin treatment. Scale bar = 1 cm

**Supplementary Figure 11.**
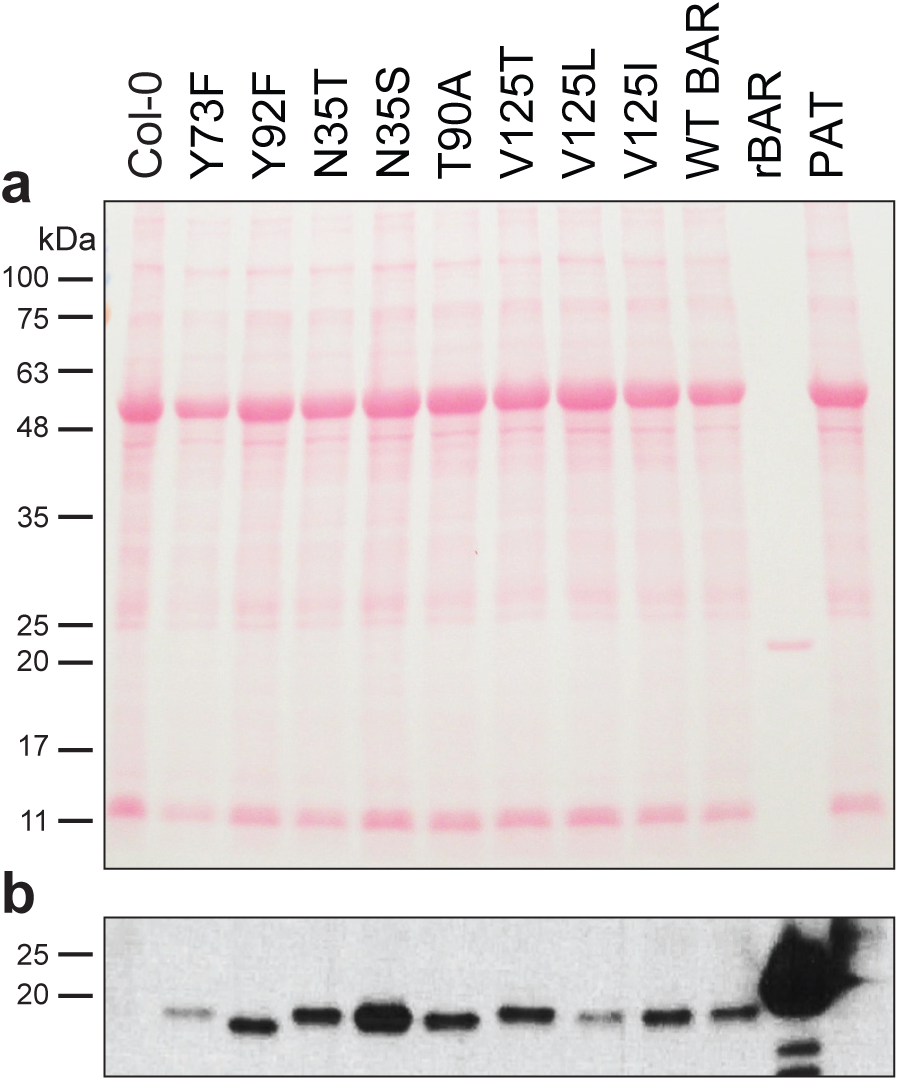
Protein levels of BAR variants in Arabidopsis. (**a**) Total proteins were extracted from T2 plants, separated by SDS-PAGE, transferred to nitrocellulose membrane and stained by Ponceau S. For each protein extraction, equal amounts of aerial tissues from 5-6 T2 populations grown from seeds from independent T1 plants were pooled. (**b**) Detection of BAR by anti-BAR immunoblotting of the membrane shown in panel (**a**). rBAR, recombinant BAR from *E.coli*.

**Supplementary Figure 12.**
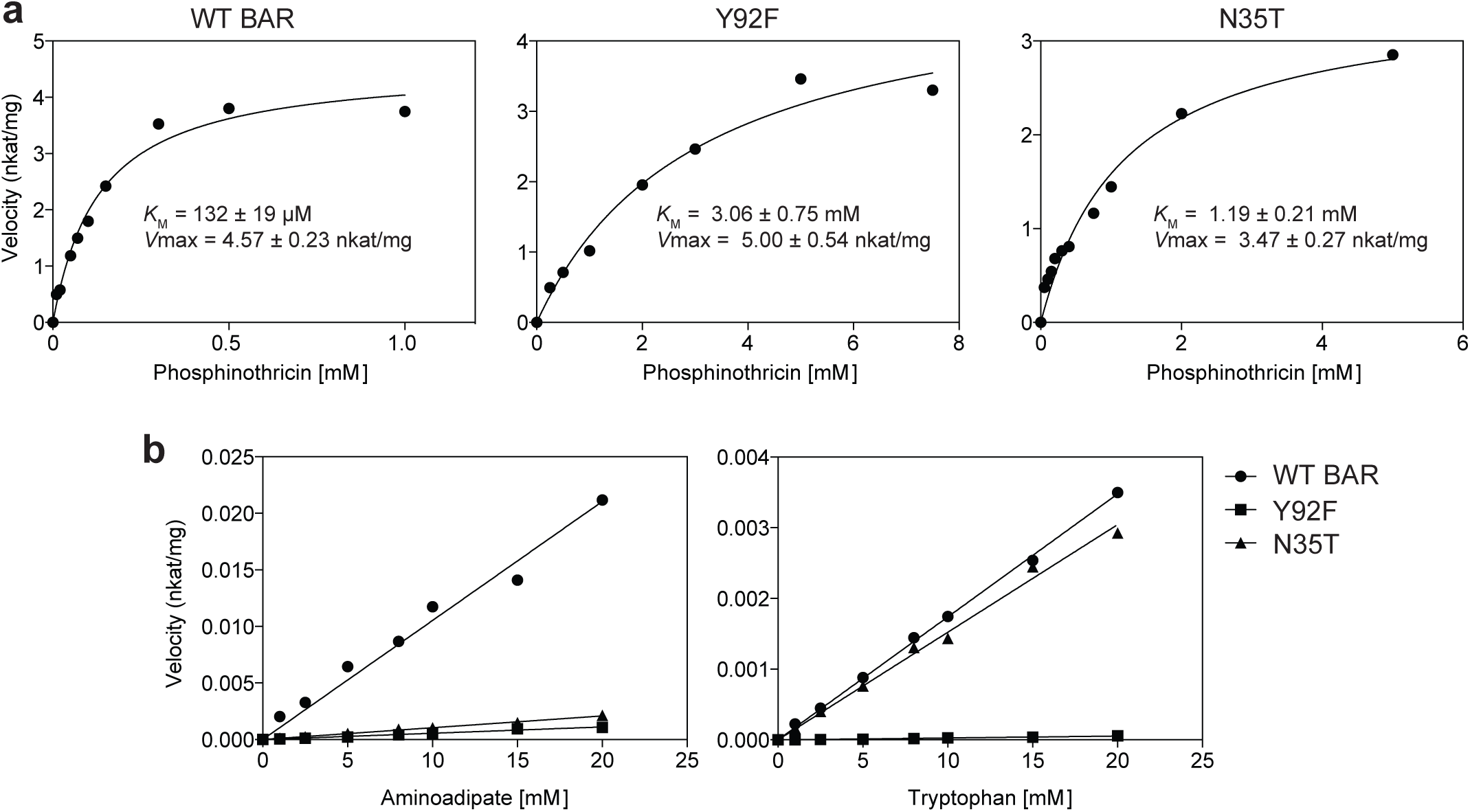
In vitro enzyme kinetic assays of wild-type BAR (WT BAR, as shown in Fig. 3) and BAR variants Y92F and N35T against native (**a**) and non-native substrates (**b**). (**a**) Calculated *K*_M_ and *V*max values for phosphinothricin are indicated. (**b**) Note that aminoadipate and tryptophan reached solubility limit before reaching saturation concentration for WT BAR, Y92F and N35T.

